# Structure of the ancestral TRPY1 channel from *Saccharomyces cerevisiae* reveals mechanisms of modulation by lipids and calcium

**DOI:** 10.1101/2020.10.12.336495

**Authors:** Tofayel Ahmed, Collin R. Nisler, Edwin C. Fluck, Marcos Sotomayor, Vera Y. Moiseenkova-Bell

## Abstract

Transient Receptor Potential (TRP) channels have evolved in eukaryotes to control various cellular functions in response to a wide variety of chemical and physical stimuli. This large and diverse family of channels emerged in fungi as mechanosensitive osmoregulators. The *Saccharomyces cerevisiae* vacuolar TRP yeast 1 (TRPY1) is the most studied TRP channel from fungi, but the molecular details of channel modulation remain elusive so far. Here, we describe the full-length cryo-electron microscopy structure of TRPY1 at 3.1 Å resolution. The structure reveals a distinctive architecture for TRPY1 among all eukaryotic TRP channels with an evolutionarily conserved and archetypical transmembrane domain, but distinct structural folds for the cytosolic N- and C-termini. We identified the inhibitory phosphatidylinositol 3-phosphate (PI(3)P) lipid binding site, which sheds light into the lipid modulation of TRPY1 in the vacuolar membrane. The structure also exhibited two Ca^2+^-binding sites: one in the cytosolic side, implicated in channel activation, and the other in the vacuolar lumen side, involved in channel inhibition. These findings, together with data from molecular dynamics simulations, provide structural insights into the basis of TRPY1 channel modulation by lipids and Ca^2+^.

## INTRODUCTION

Osmoregulation is the ability of cells to detect and counterbalance osmotic concentration variations in their surroundings^1^. In their natural environment, unicellular eukaryotic organisms like yeast, which live on plants and animals, can experience rapid water efflux (hyperosmotic shock) leading to shrinkage, or water influx (hypoosmotic shock) causing them to swell^1^. To survive hyperosmotic shocks, yeasts have been shown to release Ca^2+^ from intracellular stores to initiate a defense response that counterbalances rapid changes of osmotic concentration in their environment^2^.

The vacuole acts as a Ca^2+^ buffering system that maintains low free cytosolic Ca^2+^ concentration in yeast^3^. It has been reported that Ca^2+^ concentration in the yeast vacuole is around 1.3 mM, while cytosolic Ca^2+^ is only 260 nM^3–5^. The *Saccharomyces cerevisiae* vacuolar transient receptor potential yeast 1 (TRPY1, also known as yeast vacuolar conductance 1 or YVC1), considered the closest extant representative of the ancestral state of eukaryotic TRP channels, has been shown to play a major role in Ca^2+^ release from the vacuole to the cytosol in response to hyperosmotic stress^5^. TRPY1 is a non-selective Ca^2+^ permeable and polymodal cation channel, like many other TRP channels^6–9^. It was originally characterized as a Ca^2+^-activated channel, but was later proposed to be also mechanosensitive^10–12^. Increased cytosolic Ca^2+^ concentration can enhance TRPY1 mechanosensitivity, consistent with its proposed *in vivo* function in osmotic regulation^13,14^. Recently, it was shown that a high concentration of Ca^2+^ in the vacuolar lumen inhibits TRPY1 through Ca^2+^-binding at the S5-S6 linker^15^. TRPY1 can be also activated by reducing agents and inhibited by phosphatidylinositol 3-phosphate (PI(3)P), suggesting that reactive oxygen species and PI(3)P lipid are endogenous modulators of the channel^14^.

A recent comprehensive study on TRP channel mechanosensitivity has shown that the majority of mammalian TRP channels are insensitive to membrane stretch^16^. Nevertheless, a *Drosophila* TRP channel known as NOMPC has been shown unequivocally to be mechanosensitive both in *vitro* and in *vivo*^17^. Electrophysiological experiments suggest that TRPY1 is responsive to membrane stretch, indicating that both channels are mechanosensitive, although their activation mechanisms might be different^13,18^. Recently, a NOMPC structure revealed how its large ankyrin-repeat domain organizes into a spring-like bundle that would interact with the cytoskeleton to control channel opening^19^. In contrast, TRPY1 does not have ankyrin-repeats, and its activation might be triggered by membrane stretch sensed by its transmembrane core or by Ca^2+^ binding to its cytosolic domain^13,20^.

To further understand the structural topology of this ancestral eukaryotic TRP channel and to shed light on mechanisms of TRPY1 channel modulation by Ca^2+^ and membrane lipids, we determined the structure of the full-length TRPY1 channel in the presence of Ca^2+^ by cryo-electron microscopy (cryo-EM). The transmembrane core of the channel revealed a “classic” TRP channel domain-swapped topology, but a different arrangement of N- and C-terminal domains contributed to a distinct overall TRPY1 channel architecture. This structure of the TRPY1 channel has also revealed key PI(3)P and Ca^2+^-binding sites. Insights from all-atom molecular dynamics (MD) simulations confirmed that PI(3)P and vacuolar Ca^2+^ play a central role in maintaining a closed state of the channel, while cytosolic Ca^2+^ is important for the structural integrity of the TRPY1 cytosolic domain. In addition, our findings provide structural insights that might guide the exploration of the molecular basis of TRPY1 channel modulation by hyperosmotic stress.

## RESULTS

We overexpressed the full-length *Saccharomyces cerevisiae* TRPY1 channel in *Saccharomyces cerevisiae* and purified the channel using digitonin and glyco-diosgenin detergents. Since TRPY1 is activated by cytosolic Ca^2+^ between 10 μM to 1 mM and inhibited at 1 mM Ca^2+^ from the vacuolar luminal side^15^, we determined TRPY1 structures in an apo condition at 3.0 Å and in a 2 mM Ca^2+^-supplemented condition at 3.1 Å resolution by cryo-EM (Extended Data Figs. 1–2). Both of these cryo-EM density maps were of sufficient quality for *de novo* model building for the majority of the TRPY1 channel (Extended Data Figs. 1–4). These structures are nearly indistinguishable from each other (RMSD 0.307), including the bound Ca^2+^ ions and PI(3)P molecules (Extended Data Figs. 1–2). While Ca^2+^ was not added to the buffer solutions for the structure determined at the apo condition, there was still sufficient Ca^2+^ present that the channel was trapped in an apparently saturated Ca^2+^-bound state. This may have been due to residual Ca^2+^ leeching from the filter paper during grid preparation, a possibility that has been reported in a separate cryo-EM study of inositol triphosphate receptors^21^. As both TRPY1 structures appear to be in identical closed states, we will only discuss the 2 mM Ca^2+^-supplemented state in this manuscript (Fig. 1, Extended Data Fig. 1–4, Table 1).

**Figure 1:**
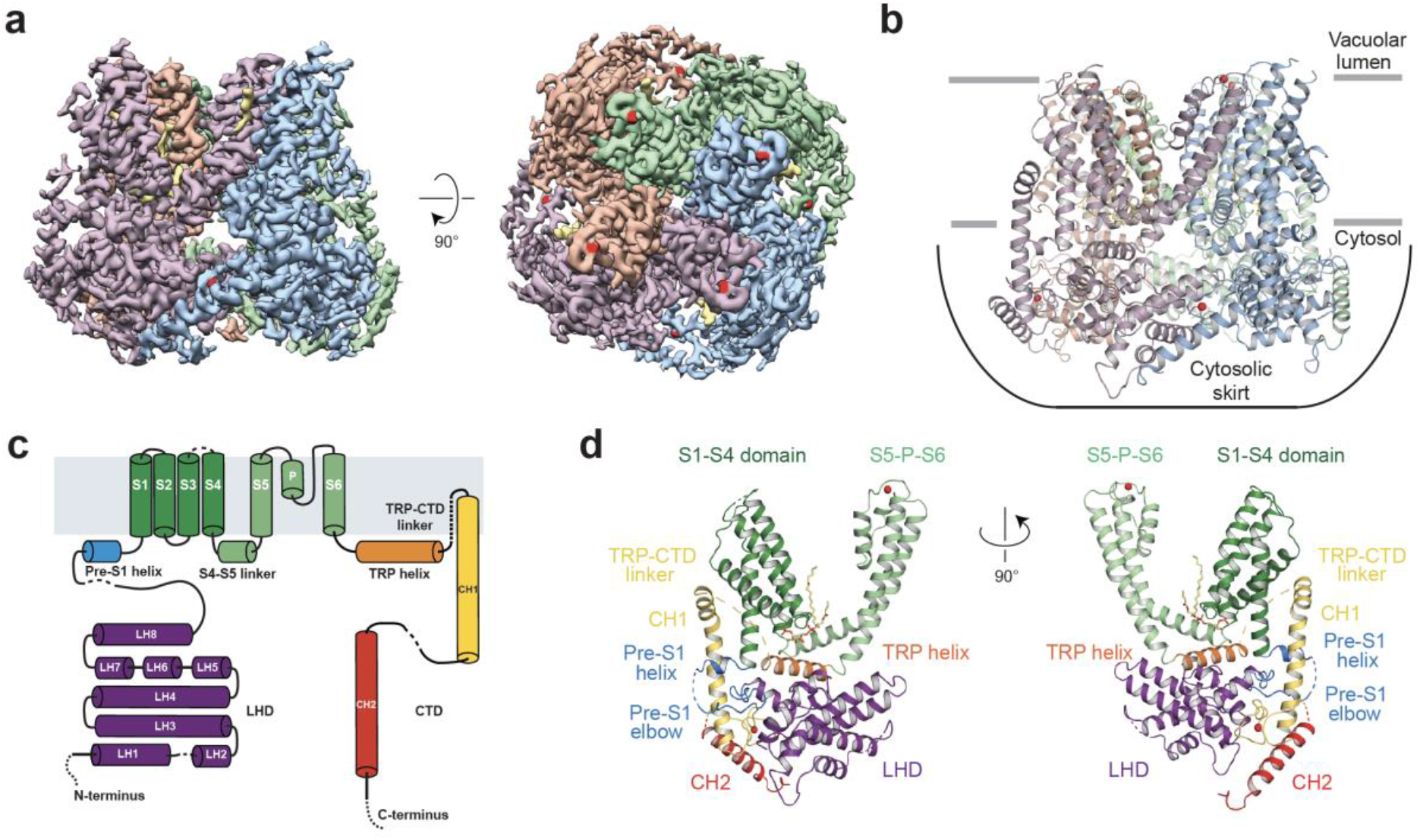
Overall architecture. **a,** TRPY1 homo-tetramer density at 3.1 Å resolution, shown at threshold level 0.031 in Chimera^62^. Side view parallel to the membrane (left) and view from the vacuolar lumen side (right) are displayed to show domain-swapped architecture and positions of lipids (khaki) and Ca^2+^ (red). **b,** Cartoon representation of the atomic model built in the cryo-EM density. Due to ambiguity, models for the lipid (khaki) at the top right corner in (a, left) were not built. Due to the absence of ankyrin-repeats, a cytosolic skirt is formed by the distinctive arrangement of the different cytosolic domains, loops and parts of helices, as labelled. Cartoon colors are matched to the electron density in **(a)**. **c,** Schematic domain organization of a TRPY1 subunit. Dashed lines indicate unmodeled traces in the flexible regions containing weak density. Helices and loops are not drawn to scale. **d,** Cartoon representation of the domain organization in a TRPY1 subunit to show relative positions of different domains, using matching colors from **(c)**.

**Table 1.** Refinement statistics for Ca^2+^-bound TRPY1 map (EMDB: 21672) and model (PDB: 6WHG):

|  |  |
| --- | --- |
| Data collection and processing | TRPY1, 2mM Ca<sup>2+</sup> condition |
| Magnification | 81,000x |
| Detector mode | super-resolution |
| Voltage (kV) | 300 |
| Defocus range (μm) | 0.5-2.5 |
| Pixel size (Å) | 1.06 |
| Total extracted particles (no.) | 3,120,706 |
| Refined particles (no.) | 254,509 |
| Final particles (no.) | 55,593 |
| Symmetry imposed | C4 |
| Map sharpening *B* factor (Å<sup>2</sup>) | -94.37 |
| Map resolution (Å) | 3.1 |
| FSC threshold | 0.143 |
| **Model Refinement** |  |
| Model resolution cut-off (Å) | 3.1 |
| Model composition |  |
| Nonhydrogen atoms | 17116 |
| Protein residues | 2092 |
| Ligands | PWE: 4, CA: 8 |
| R.M.S. deviations |  |
| Bond Lengths (Å) | 0.009 |
| Bond angles (°) | 1.129 |
| Validation |  |
| Molprobity score | 1.47 |
| Clashscore | 2.65 |
| Poor rotamers (%) | 1.98 |
| CαBLAM outliers (%) | 1.60 |
| EMRinger score | 3.24 |
| Ramachandran plot |  |
| Favored (%) | 96.87 |
| Allowed (%) | 3.13 |
| Disallowed (%) | 0.00 |

Similar to previously determined TRP channel structures^22–25^, TRPY1 forms a domain-swapped homo-tetramer with each subunit having a cytosolic N-terminal linker helical domain (LHD) consisting of eight tightly packed α-helices, six transmembrane α-helices (S1-S6) with a pore helix (P helix), followed by a TRP helix and a C-terminal domain (CTD) mainly comprising two long α-helices (Fig. 1). Due to the absence of the ankyrin-repeat domains at the N-terminus (Extended Data Fig. 5), the entire cytosolic domain spanning LHDs and CTDs across the four TRPY1 subunits assumes the shape of a cytosolic “skirt” (Fig. 1b), discussed in detail along with computational models in later sections. Although we worked with the full-length TRPY1 protein construct (Met1-Glu675), parts of the N-terminal domain (residues Met1-Asn24, Asp55-Glu65), the pre-S1 elbow (residues Leu216-Phe225), the loop between S3 and S4 helices (residues Pro323-Lys327), the TRP-CTD linker (residues Ala487-Ser529), and some parts of the C-terminal domain (residues Leu572-Ser580, Asp608-Glu675) were not resolved in our structure likely due to flexibility (Fig. 1) and models were not built for them. We also did not build models for other ambiguous densities in the transmembrane region that are most probably annular lipids or detergent molecules. Overall, the TRPY1 structure has several “classical” TRP channel domains seen in other TRP channel structures (Fig. 1c, d), yet there are specific features relevant for its regulation by Ca^2+^ and the channel’s role in mechanosensation in the yeast vacuolar membrane (Fig. 1).

In our TRPY1 structure, the LHDs have distinct features in terms of the number and length of the helices, their relative arrangement, and the length of LH5-LH6 loop (Fig. 1c, Fig. 2a and Extended Data Fig. 6a). Structural comparison with counterparts from NOMPC (PDB: 5VKQ)^19^, TRPM4 (PDB: 5WP6)^26^ and TRPC6 (PDB: 5YX9)^27^ reveals good conservation for the five LHD helices LH4-LH8 that are located adjacent to the pre-S1 helix (Fig. 1c, d and Extended Data Fig. 6b). However, LH3 in TRPY1 is two helical turns longer than in NOMPC, TRPM4 and TRPC6 (Extended Data Fig. 6b). Toward the N-terminus, like TRPM4, TRPY1 LHD has two more helices, LH1 and LH2. This is in contrast with NOMPC and TRPC6 where three such helices are present at the N-terminus, two of which are very short while the other is similar in length to LH2 of TRPY1. The LH5-LH6 loop (Arg154-Asn169) in TRPY1 is comparable to TRPC6 but longer than in NOMPC and TRPM4 (Extended Data Fig. 6b). Strikingly, the LH5-LH6 loop in TRPY1 interacts with the CTD of the neighboring subunit (discussed later) (Fig. 2a and Extended Data Fig. 6a). This loop-mediated interaction is a distinct feature of TRPY1’s cytosolic skirt and is not observed in other TRP channels. It is remarkable that such a lengthy loop remained stable at this position and consequentially a high-quality electron density was captured to model side chains for the entirety of the loop. The reason for this stability can be partly attributed to several interactions within the LHD such as three salt bridges between Arg155 and Glu172, Lys102 and Glu160, and Arg33 and Asp162 (Fig. 2a). Coordination of a bridging cytosolic Ca^2+^ at the distal end of the LH5-LH6 loop provides additional stability (Fig. 2a). In the absence of the ankyrin-repeat domains in TRPY1, the cytosolic skirt, especially the LHDs may harbor the docking site for binding partners in the *S. cerevisiae* cytosol. Therefore, the structural differences in LHD helices along with the longer LH5-LH6 loop and its capability to coordinate a Ca^2+^ ion might have implications for TRPY1 channel function.

**Figure 2:**
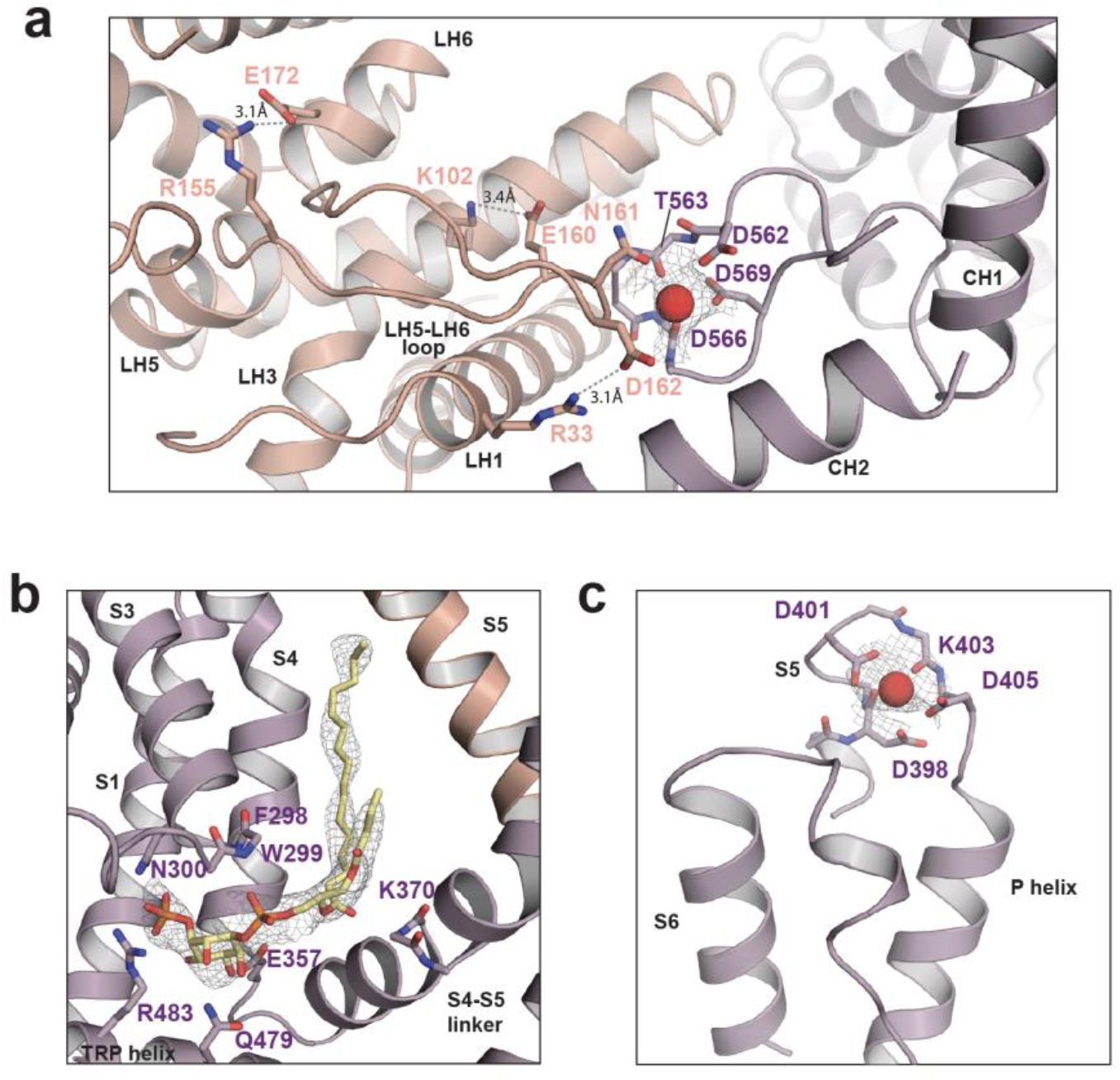
Ca^2+^ and PI(3)P binding sites. **a,** Cytosolic Ca^2+^-binding site. Ca^2+^ is displayed in red. Interacting protein residues are displayed in sticks and labelled accordingly. **b**, PI(3)P binding site. The atomic model of PI(3)P lipid is displayed in sticks with elements colored as: carbon in khaki, phosphorous in orange and oxygen in red. **c,** Ca^2+^-binding sites at the vacuolar lumen side. Shown as in (a). Densities for the cytosolic Ca^2+^, PI(3)P and luminal Ca^2+^ are isolated from the original map and displayed in grey meshes at sigma level 1.5 in (a), (b) and (c), respectively.

Another unique feature of the TRPY1 structure is the position of the CTD relative to the vacuolar membrane and the interaction of the CTD with the N-terminal LHD of a neighboring subunit (Fig. 1 and Extended Data Fig. 6a). The CTD is composed of a TRP-CTD linker that is not resolved in our cryo-EM structure, two long α-helices labeled CH1 and CH2, and a connecting CH1-CH2 loop (Fig. 1). The CH1 helix is partially embedded in the membrane (Fig. 1b) and connected to the CH2 helix by a loop that harbors the cytosolic Ca^2+^-binding site (Fig. 1b, c and Fig. 2a). This cytosolic Ca^2+^ is coordinated by backbone and sidechain oxygen atoms from Asp562, Thr563, Asp566 and Asp569 on the CH1-CH2 loop, as well as from Asn161 on the LH5-LH6 loop of the neighboring LHD (Fig. 2a). Previously, it was suggested that the acidic patch of the four tandem aspartates between Asp573-Asp576 comprises the cytosolic Ca^2+^ binding site in TRPY1^13^ (Extended Data Fig. 6a and 7). Because we could not resolve the density for the seven amino acids between Asp574-Ser580 (Extended Data Fig. 6a and 7) due to flexibility, it is not conclusive from our structure whether this acidic patch indeed is capable of binding Ca^2+^. However, it is conceivable that, if a Ca^2+^ ion were to bind at this patch in our purified protein, then it would lead to a greater stabilization of the CH1-CH2 loop allowing us to elucidate the complete structure of the same loop. Moreover, a recent mutagenesis study has reported that this acidic patch is not essential for Ca^2+^ activation of TRPY1^14^. Furthermore, the acidic patch is not conserved as revealed by the sequence alignment from multiple fungal genomes (Extended Data Fig. 7). Therefore, it is unlikely that this acidic patch binds Ca^2+^ in purified TRPY1 or in *S. cerevisiae*. Rather, TRPY1 activation by cytosolic Ca^2+^ is mediated by Ca^2+^ binding at the subunit interface as discussed above. Interestingly, Asn161 at the subunit interface is conserved among three (*S. cerevisiae* TRPY1, *K. lactis* TRPY2, *C. albicans* TRPY3) of the six sequences aligned across different fungal species (Extended Data Fig. 7). In the other three (*N. crassa, A. niger, A. flavus*), an acidic aspartate variation has replaced this asparagine and the CH1-CH2 loop (Arg553-Asp581) is shortened by 6-7 amino acids (Extended Data Fig. 7). The implications of this sequence variation and loop shortening in these three fungal genomes remain unknown, but our analyses suggest the presence of a similar cytosolic skirt structure and cytosolic Ca^2+^-induced activation mechanism for at least TRPY1, TRPY2 and TRPY3.

The transmembrane domain of the channel was resolved to around 2.8 Å resolution (Extended Data Fig. 1f), which allowed us to visualize a lipid density that we assigned to PI(3)P (Fig. 2b) and a small non-protein density that we assigned to a luminal Ca^2+^ (Fig. 2c). The PI(3)P lipid was co-purified with the channel from yeast, as lipids were not added during purification. Endogenous yeast PI(3)P is typically found on the intraluminal vesicles of endosomes and vacuoles^28^, and it has been shown to inhibit TRPY1 channel activity^14^. The PI(3)P binding site is located between the S1-S4 domain, the S4-S5 linker and the TRP helix of one subunit and S5 of an adjacent subunit (Fig. 2b). The inositol ring of PI(3)P is positioned underneath the S1-S4 domain and is wedged above the TRP helix. In this position, the inositol ring interacts with TRP helix residues Arg483 and Gln479 via its 3’ phosphate and 2’ hydroxyl groups, respectively (Fig. 2b). The PI(3)P position in this pocket is further stabilized by hydrogen bonding between Glu357 and the 5’ hydroxyl on the inositol ring as well as between the backbone amine of Lys370 and the carboxyl from the ester moiety on one of the acyl tails of the lipid (Fig. 2b). Apart from this, the 1’ phosphate of PI(3)P maintains hydrogen bonding with the backbone amines of Phe298 and Trp299 and thus locks the hydrophilic headgroup of PI(3)P in position. The two acyl chains of PI(3)P branch out in a ‘V’ shape. One tail stabilizes in a cleft formed by S4 and the S4-S5 linker of one subunit and S5 and S6 of an adjacent subunit, through hydrophobic interactions. The other tail resides near the S4-S5 linker and we did not capture most of its density probably due to flexibility. A similar position for an inhibitory phosphatidylinositol lipid was observed in the TRPV1 structure^23^, suggesting a conserved mechanism of lipid-mediated modulation among TRP channel homologues.

The luminal Ca^2+^-binding site is formed by the loop connecting the S5 and P helices. The bound Ca^2+^ ion is coordinated by the sidechains of residues Asp398, Asp401 and Asp405, as well as by the backbone carbonyls of Asp398 and Lys403 (Fig. 2c). Since we have captured a closed state of TRPY1, this observation agrees with recent results^15^ suggesting that two of these residues, Asp401 and Asp405, are involved in the direct binding of Ca^2+^ and are therefore critical for Ca^2+^-dependent inhibition of TRPY1. The functional importance of this Ca^2+^-binding site is further underscored by the fact that Gly402Ser mutation resulted in a constitutively active channel^29^. However, how the luminal Ca^2+^ leads to rearrangement of the S6 helix to stabilize the closed conformation of the channel remains to be elucidated. The removal of the luminal Ca^2+^ would render the P helix flexible, transmitting a gating signal to the pore in S6. We also observe some additional features in TRPY1 such as a π-hinge in the middle of S6, similar to some other TRP channels and two proline residues, Pro432 and Pro433, at the beginning of S6 (Extended Data Fig. 7 and 8). In the absence of luminal Ca^2+^, these features might have implications in channel gating by allowing more flexibility in S6.

It is interesting to note that the TRPY1 pore structure is almost identical to the NOMPC pore in the closed state, and similar to the pore structures from TRPA1, TRPM4 and TRPC6 (Fig. 3 and Extended Data Fig. 8). Like all other TRP channels, the TRPY1 pore is comprised of a selectivity filter and a lower gate (Fig. 3b). The selectivity filter is formed by the backbone carbonyl oxygen atoms of residues Leu419 and Gly420 (Fig. 3a). Just above the selectivity filter, two negatively charged residues, Asp425 and Glu428, could attract and interact with luminal cations entering the pore. Overall, the luminal side of the pore is strongly negatively charged (Extended Data Fig. 9) as the luminal Ca^2+^ binding site residues such as Asp398, Asp401 and Asp405 also line the surface (Fig. 2c and 3a). The diameter of the selectivity filter is around 5.6 Å at the Gly420 carbonyl and is large enough to accommodate partially hydrated cations. This is similar to other TRP channels solved in the open state^22,30^. Therefore, although the selectivity filter is open in TRPY1, the pore is still closed to ion permeation due to a hydrophobic seal at the lower gate formed by Ile455, where the pore diameter is constricted to 1.4 Å (Fig. 3b).

**Figure 3:**
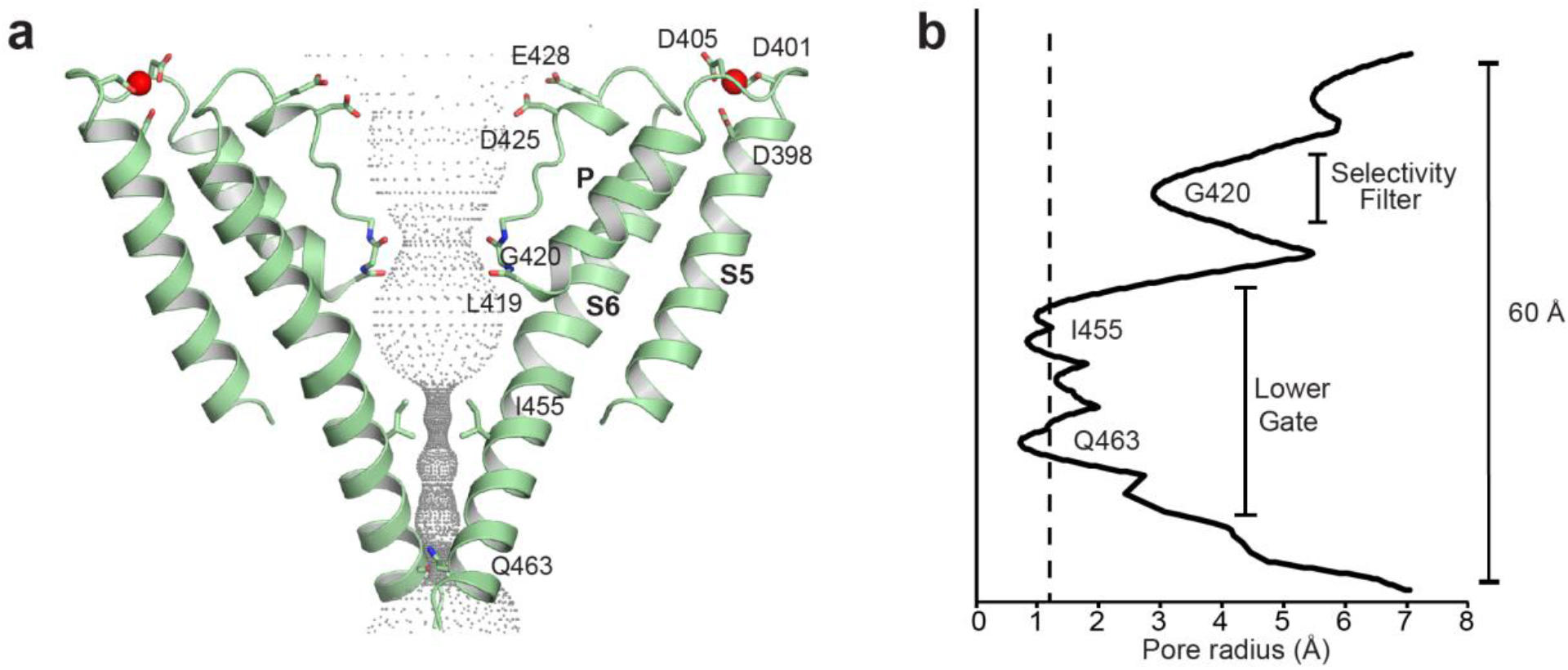
Architecture of the ion permeation pathway. **a,** Cartoon representation of TRPY1 ion permeation pathway. Ca^2+^ is displayed in red. Grey dots illustrate pore size as determined by HOLE. **b,** Graphical representation of the radius of the TRPY1 pore as a function of the distance along the ion permeation pathway. The dashed line indicates the radius of a dehydrated Ca^2+^.

Another interesting aspect of the TRPY1 structure is the network of interactions within each of the subunits that we identify as crucial in stabilizing the closed state of the channel (Fig. 4). These interactions involve the TRP helix, the S4-S5 linker, the LHD, and the CH1-CH2 loop (Fig. 4). The S4-S5 linker forms multiple contacts with the TRP helix which includes a salt bridge interaction between the side chains of Arg360 and Asp470, multiple hydrogen bonds between the side chain hydroxyl of Tyr473 and backbone amide of Arg360, and backbone carbonyls of Glu357 and Ser358 (Fig. 4a). Evidently, Tyr473 is centrally positioned among these residues and the majority of its interacting partners are backbone atoms from the S4-S5 linker. Tyr473 is also conserved across the fungal genomes that we aligned (Extended Data Fig. 7). Together, these observations suggest that Tyr473 is a pivotal residue in the stabilization of the closed state of TRPY1. Earlier studies showed that mutations Tyr458His and Tyr473His increase the open probability of the channel^29^. These two mutants caused unstable open and closed states of the channel, suggesting their involvement in channel gating^29^. As deducible from our structure, the first mutant likely destabilizes the S6 helix with either a direct effect on the pore or an indirect effect on the stability of the Arg360-Asp470 salt bridge, or both. The second mutant directly disrupts the three hydrogen bonding interactions between the TRP helix and the backbone of the S4-S5 linker. Intriguingly, PI(3)P is positioned right next to the TRP helix and the S4-S5 linker and provides additional stability to this region, as discussed earlier (Fig. 2b). Therefore, it is possible that PI(3)P interaction and channel gating is coupled in TRPY1. Finally, two salt bridges that are worth mentioning are formed within the same subunit between Arg190 and Arg197 from the LH8 of LHD with Asp557 and Glu560 from the CH1-CH2 loop of the CTD, respectively (Fig. 4b). Together, they seem to stabilize the communication between LHD and CTD in each of the subunits. Overall, an extensive network of salt bridges and hydrogen bonding between different domains of TRPY1 play an important role to maintain the closed state of the channel.

**Figure 4:**
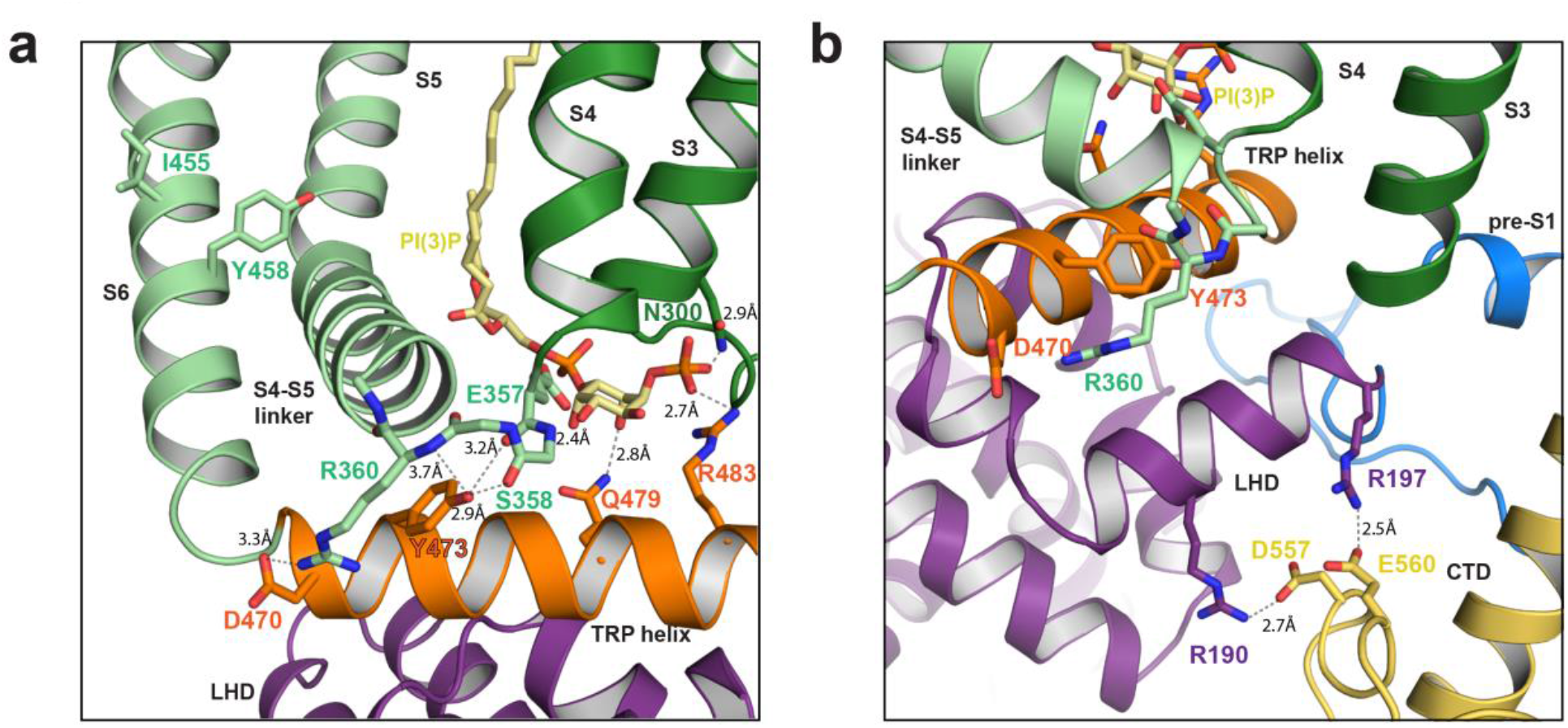
Interaction network in the closed state of TRPY1. **a,** Interaction network between S4-S5 linker (green), TRP helix (orange) and PI(3)P (khaki) to stabilize the closed state of TRPY1. **b,** Interactions between LHD and CTD.

To further elucidate the roles of PI(3)P and Ca^2+^ in the stabilization of the TRPY1 closed conformation, all-atom MD simulations were carried out on the TRPY1 structure. Five systems were built and simulated (Extended Data Table 1), each representing a different potential state that the channel may exist *in vivo* or *in vitro*. Four separate systems (Sim1 to Sim4 in Extended Data Table 1) had the TRP-CTD linker (Ala487-Ser529) in a “vertical” configuration (Extended Data Fig. 10). These include All-Bound (Sim1), in which all PI(3)P lipids and Ca^2+^ ions are present; None-Bound (Sim2), in which all have been removed; No-Inh (Sim3), in which only the cytosolic Ca^2+^ sites are occupied while luminal Ca^2+^ ions and PI(3)P lipids are removed; and the Y473H model (Sim4), in which this gain of function mutation^31^ (see “network of interactions within each of the subunits” above) was introduced to the No-Inh system in all four subunits. A fifth system featured the TRP-CTD linker in a “horizontal” configuration (Extended Data Fig. 10) with all PI(3)P lipids and Ca^2+^ ions present. Each system was prepared and simulated with an identical protocol that included 100 ns of production simulations.

The All-Bound horizontal and vertical configurations exhibited noticeable differences in water permeation across the lipid bilayer during simulation (Extended Data Fig. 11). When averaged over the entire simulation trajectory, some water density was observed within the lipid bilayer near the TRPY1 S1-S4 helices and the TRP-CTD linker in both systems, but to a far greater extent in the vertical configuration compared to the horizontal configuration (Extended Data Fig. 11a, c, d and f). There is precedence for the facilitation of ion permeation across the lipid bilayer by similar domains from potassium channels, as the isolated voltage-sensing domain (VSD) of the Shaker Kv channel has been shown to form a cation selective pore^32^. Voltage-dependent permeation of protons was also shown for the ion channel HV1^33,34^, which has only four transmembrane helices in a VSD-like arrangement (S1-S4) with a predicted hydrated pathway^33^. The individual S1-S4 subunits of TRPY1 coupled to the TRP-CTD linker and the CH1 helix featuring several charged residues at its N-terminus (Lys514 to Asp518), may function in a similar manner, explaining the observed water permeation. Although both vertical and horizontal configurations are feasible, we focused all subsequent analyses and discussion on our simulations of the channel with the TRP-CTD linker in the vertical configuration.

Simulations revealed that removal of PI(3)P resulted in early indications of pore opening and an overall increase in the dynamics of the TRPY1 structure. In the None-Bound, No-Inh, and Y473H systems, the S4 helix exhibits a greater degree of displacement from its initial position compared to the All-Bound system (Extended Data Fig. 12a-c). A similar result is seen for the S4-S5 linker (Extended Data Fig. 12d-f). The removal of PI(3)P facilitates a greater displacement of key helices likely involved in gating and that may lead to opening of the pore at longer timescales (Extended Data Fig. 12a-f). Additionally, both helices (S4 and S4-S5 linker) in the Y473H system exhibited the greatest degree of deviation, indicating that this mutation influences the dynamics of the transmembrane helices. Removal of PI(3)P also had an apparent effect on the dynamics of the TRP helix, as the distance between opposing TRP helices between two subunits maintained a smaller separation in the All-Bound system compared to the other three simulated systems (Extended Data Fig. 13g-i). These events may facilitate opening of the pore when PI(3)P is removed. Finally, removal of PI(3)P disrupted networks of allosteric communication between the different domains within the same subunit of TRPY1, as revealed by a dynamical network analysis^35^ (Extended Data Fig. 14). In the All-Bound system, the optimal path of allosteric communication from the N-terminus to the end of the S4-S5 linker (Gly26-Ser376) goes directly from the TRP helix to the S4-S5 linker through Tyr473 (Extended Data Fig. 14a, b), while in every other simulated system the optimal path does not pass through Tyr473 (Extended Data Fig. 14e-f, i-j, and m-n), indicating that the presence of the PI(3)P lipid maintains a more compact network of communication between the TRP helix and the S4-S5 linker. Similarly, the optimal path from the end of the S4-S5 linker to Ile291 at the base of the S2 helix first passes through the S3 and the S2 helix in the All-Bound system (Extended Data Fig. 14c, d). In the other three systems, in which PI(3)P is not present, the optimal path does not pass through the S2-S3 helices (Extended Data Fig. 14g-h, k-l, and o-p). All our analyses suggest that PI(3)P stabilized the closed state.

Analyses of interactions required for the stabilization of the closed state revealed that the Arg197-Glu560 salt-bridge was disrupted to a greater degree in simulations of the None-Bound and No-Inh systems relative to the All-Bound system (Extended Data Fig. 13a-c). Other interactions required for the stabilization of the closed state, such as Arg360-Tyr473 and Lys193-Tyr473, were disrupted by mobile lipid molecules from the bilayer (Extended Data Fig. 13d-f). As these lipid molecules formed interactions with one of the residues involved, the native interaction was disrupted, demonstrating the importance of the surrounding lipid environment to the function of TRPY1.

Next, ion density was analyzed by averaging the location of all K^+^ ions over the 100-ns long trajectories for all systems. Significant differences between the All-Bound and No-Inh systems were recorded (Extended Data Fig. 15a-d). In contrast to the All-Bound system, K^+^ density was observed beyond the selectivity filter formed by Leu419 and Gly420 in the No-Inh system (~95-100 Å in the *z*-axis; Extended Data Fig. 15c, d). This indicates that the removal of bound luminal Ca^2+^ and more mobility of the associated loops, removal of PI(3)P and greater displacement of transmembrane helices, or a combination of both resulted in the positively charged species having an increased accessibility to the pore of TRPY1, in support of the experimental observations indicating that luminal Ca^2+^ inhibits opening of the channel^15^.

As with the luminal Ca^2+^-binding sites, Ca^2+^ binding to the cytosolic sites also affected the dynamics and permeation properties of the channel. The Val50-Ser68 loop forms the neck of the cytosolic skirt that may modulate cytosolic ion permeability. The dynamics of this domain depend on the presence of Ca^2+^ as revealed by simulations. In the None-Bound system, the distance between the center of mass of these loops in opposing subunits was smaller than for the other three systems, and the N-terminal domain exhibited greater deviation from its starting position, indicating that Ca^2+^ “locks” these domains in place (Extended Data Fig. 13j-l, 12g-i, respectively). This is also revealed in the analysis of average pore radius over the entire trajectory (Fig. 5). Closing of this cytosolic skirt constricts the pore radius within the skirt in the None-Bound system (Fig. 5f), and provides a mechanistic explanation for how the presence of cytosolic Ca^2+^ leads to a higher open probability observed in experiments^14,15^. In addition, the density of K^+^ varies in the cytosolic domain depending on the presence or absence of Ca^2+^. In all systems where cytosolic Ca^2+^ is bound, there is an apparent concentration of K^+^ density in the 40-60 Å range in the *z*-axis (Extended Data Fig. 15a, c, e, and g). In the None-Bound system, this region is mostly devoid of K^+^ density, indicating that the closing of the cytosolic skirt as described above precludes K^+^ entry in this region.

**Figure 5:**
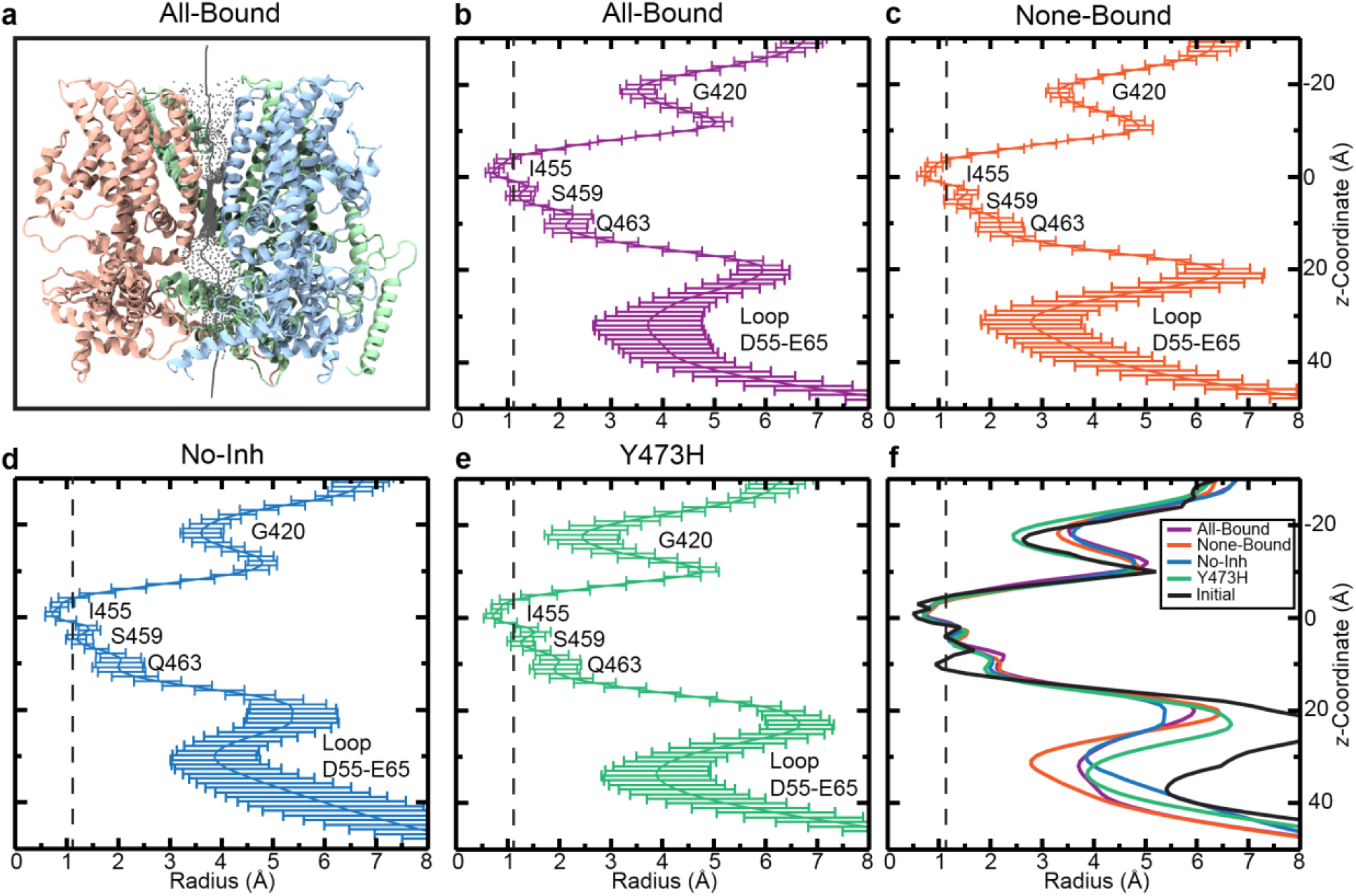
Simulations predict cytosolic Ca^2+^ binding leads to stabilization of the cytosolic skirt. **a**, Pore size through the entire channel illustrated by grey dots, and the optimal path shown by a grey line. For ease of viewing, chain A has been omitted. **b-e,** Pore radius averaged over the entire 100 ns trajectory for each system, with standard deviations for each value shown at the associated location in the *z*-axis. Important residues or motifs are indicated, and a dashed line marks the size of a dehydrated Ca^2+^ ion. **f,** The average pore radius overlaid for each system, with the initial structure (black; before simulation) that contains all modeled loops.

## DISCUSSION

TRP channels are absent in bacteria, archaea and plants, but present in all other eukaryotes including algae and fungi^36,37^. There are 28 family members in mammals that are polymodal and activated by a variety of physical stimuli and compounds. As eukaryotes evolved from single-celled to more complex multicellular organisms, TRP channel proteins may have acquired the ability to sense and respond to stimuli in more sophisticated ways. Thus, TRP channels from single-celled organisms, such as yeast, can be considered as archetypical channels providing insights into the most basic and fundamental properties common to all family members.

The *Saccharomyces cerevisiae* TRP channel homolog TRPY1 has been extensively studied over the last twenty years using yeast genetic manipulations, functional assays, and electrophysiology, yet its structure has remained elusive^14,15,31,38^. Biophysical studies have shown that TRPY1 plays a critical role in yeast Ca^2+^ homeostasis^3^. It has also been proposed that changes in Ca^2+^ concentration and mechanical force gate TRPY1^13^. Nevertheless, the molecular basis underlying this channel’s function has not been established. The high-resolution structure of TRPY1 presented here serves as a stepping-stone towards understanding TRP channels’ evolution and TRPY1 gating mechanisms modulated by Ca^2+^, lipids and membrane stretch in fungi.

Since 2003, structures for representative members from almost all mammalian TRP channel families have been determined^24,39^. These structures have revealed that mammalian TRPs share structural and functional features, which are also seen in our TRPY1 structure. The architecture of the S1-S4 domains, the pore domains comprising S5, S6 helices and the connecting pore helix, and the TRP box are all structurally conserved between TRPY1 and mammalian TRP channels, despite having low sequence similarity. This is expected since protein-folds exhibit fewer and slower changes compared to sequence during evolution^40^. Apart from the aforementioned domains, the structural similarity of TRPY1 extends further to NOMPC, TRPM4, and TRPC6 channels in terms of the overall structure of the N-terminal LHD. Like all other TRPs, TRPY1 is also a tetramer. Nevertheless, when compared to TRPM4 and TRPC6 (Extended Data Fig. 5), it is clear that TRPY1 also has some distinct structural features that might have been acquired throughout evolution or might be representative of ancestral features lost in the mammalian members. Interestingly, mammalian TRP channel structures are different compared to the TRP channel homolog in algae, TRP1^41^. This channel adopts a 2-fold symmetrical rose-shape architecture with additional structural elements not seen in TRPY1 or any of the mammalian TRP channels, suggesting that TRPY1 is more evolutionarily related to the mammalian TRP channels than TRP1^41^.

Several unique structural features of TRPY1 are likely to be relevant for channel gating. The TRP-CTD linker and the CH1 helix are tucked against the S1-S4 bundle and insert into the membrane significantly in our model (Fig. 1b, c). This arrangement provides a direct connection between the TRP helix, the periphery of the channel and the surrounding lipid bilayer, which could be relevant for mechanisms of mechanosensation involving force from lipids^42^. In addition, the CH1 helix wraps on the back of pS1 helix to connect to the cytosolic Ca^2+^-binding site and CH2, which in turn interacts with the periphery of the neighboring subunit’s LHD, a unique arrangement not seen in any other TRP channel. The stabilization of this architecture by the cytosolic Ca^2+^ that activates the channel further implicates the TRP helix in channel gating.

The presence of an endogenous PI(3)P lipid in our TRPY1 structure parallels the presence of a phosphatidylinositol lipid in a TRPV1 structure in a closed conformation, with the lipid occupying TRPV1’s vanilloid binding site^23^. The position of PI(3)P lipids in the TRPY1 structure is therefore suggestive of an active role played by lipids and compounds in channel modulation, also supporting the hypothesis of a basal and evolutionary conserved gating mechanism among TRP channels. It is conceivable that mechanical force generated by hyperosmotic shocks in yeast could induce the release of PI(3)P lipids from their binding site, thus facilitating opening of the channel and Ca^2+^ flow from the vacuole to the cytosol to start a hyperosmotic shock defense mechanism in yeast (Fig. 6).

**Figure 6:**
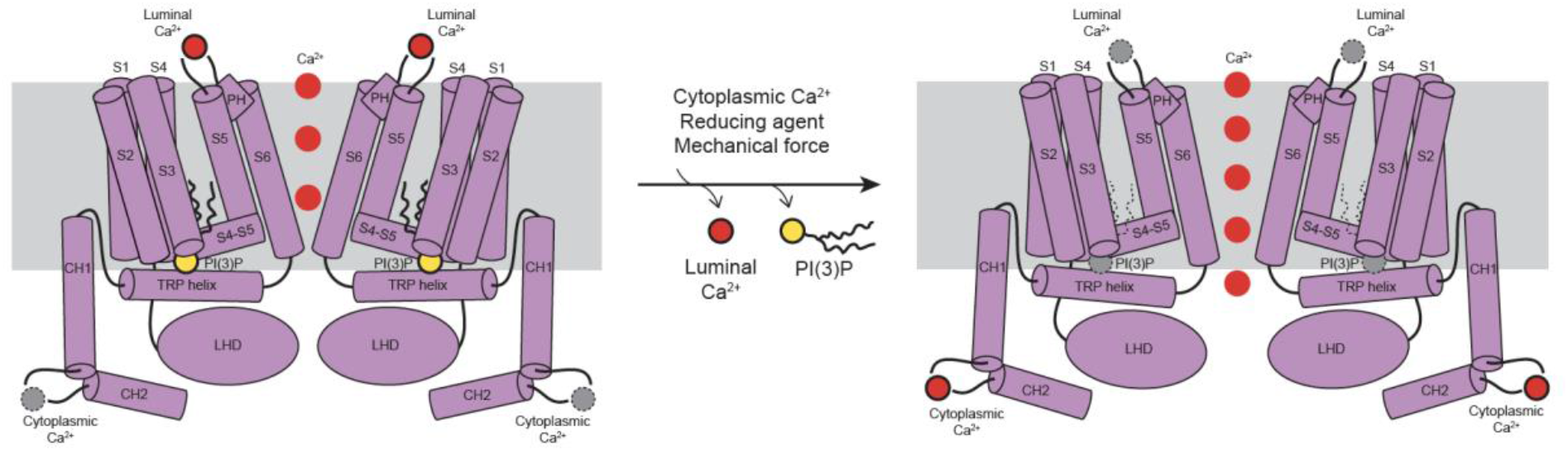
Proposed gating mechanism of TRPY1. Schematic diagram illustrating potential gating mechanism for TRPY1. On each side, two diagonal subunits of TRPY1 are shown in purple. PI(3)P is shown in khaki, and the red circles indicate the TRPY1 Ca^2+^-binding sites and the flow of Ca^2+^. For simplicity, the diagonal subunits are shown in the same topology.

The two Ca^2+^-binding sites revealed in our TRPY1 structure are distinct from those found in other TRP channels that are confined to a cavity between S2 and S3 helices, and the S2-S3 linker that resides in the membrane region^30,43,44^. Rather, in each TRPY1 subunit, one Ca^2+^ binds to the cytosolic side and another Ca^2+^ binds at the vacuolar lumen side (Fig 2a, c). It is fascinating that the two Ca^2+^-binding sites orchestrate opposite effects on the channel gating, the vacuolar lumen site being inhibiting and the cytosolic site being activating^14,15^. Simulations presented here offer insights into the possible mechanistic explanations for the experimental results^15^ concerning the role of the Ca^2+^ ions. Removal of cytosolic Ca^2+^ resulted in a domain constriction and a more occluded pore radius in this region (Fig. 5 and Extended Data Fig. 13). Removal of luminal Ca^2+^, however, led to an increase in the presence of K^+^ beyond the selectivity filter (Extended Data Fig. 15a, c), demonstrating how these unique Ca^2+^-binding sites can modulate channel activity on a molecular scale.

All sites were fully occupied by Ca^2+^ in our TRPY1 structure captured in a closed state, but it appears that only the vacuolar lumen site would remain occupied in the native yeast conditions that favor channel closure. Channel activation may require removal of Ca^2+^ from the vacuolar lumen site to create a Ca^2+^-flux directed from the vacuole to the cytosol. Yet, the concentration of Ca^2+^ in the vacuole is 3,000 times higher than in the cytosol under normal conditions^3–5^. It has been argued that vacuolar free-Ca^2+^ is scarce, as most Ca^2+^ ions are bound to vacuolar polyphosphate^45^, thus adding another layer of complexity to TRPY1 function in calcium-induced calcium release (CICR) feedback loops *in vivo*. Notably, TRPY1 is able to integrate various stress signals from the environment to produce a combined effect inside yeast cell in terms of Ca^2+^ signals. Two of the eukaryotic channels involved in CICR and studied extensively are the Inositol 1,4,5-trisphosphate and the Ryanodine receptors responsible for intracellular Camrelease in cardiac and skeletal muscles^46^. It is fascinating to note that such feedback loop to regulate an intracellular ion channel was preserved during evolution.

The TRPY1 structure reported here is in a closed state. Given that PI(3)P lipids are strongly bound to the channel, we suggest that an abundance of PI(3)P in the vacuole keeps TRPY1 in its closed conformation. The presence of PI(3)P in the observed binding site brings the TRP helix closer to the S4-S5 linker through interactions between Tyr473 and Arg360 residues. The destabilization of this interaction, which is probably the cause of gain-of-function^31^ in the TRPY1 Y473H mutant, might lead to opening of the channel upon application of different endogenous stimuli that are proposed to gate TRPY1. Our proposed model is in agreement with the notion that the S4-S5 linker is the “gear box” conveying gating force in TRP channels and suggests that TRP channels maintained a conserved gating mechanism throughout evolution. This is supported by simulations, in which the removal of PI(3)P had several effects on the dynamics and interactions implicated in channel activity. In particular, allosteric communication between the NTD and the S4-S5 linker was directly mediated by the Tyr473-Arg360 interaction only when PI(3)P was present (Extended Data Fig. 14).

The open state of TRPY1 has a conductance >300 pS (180 mM KCl), suggesting that gating requires conformational changes that can lead to a pore opening of as much as 10 Å in diameter estimated using Hille’s equation^47,48^. It is tempting to speculate that mechanical force from lipids is communicated through the TRP-CTD linker and the S1-S4 domains to the TRP and S4-S5 linker helices to gate the channel, and that this requires an intact skirt with bound Ca^2+^. However, how various parts of TRPY1 re-arrange to accommodate a larger pore that can sustain its conductance remains to be elucidated.

## MATERIALS AND METHODS

### Protein expression and purification

Wild type TRPY1 containing YepM plasmid^49^ was transfected into BJ5457 *Saccharomyces cerevisiae* (ATCC) for constitutive expression of the protein. The cells were grown at 30°C until OD600 reached mid log phase (1.2-1.4) and thereafter harvested to store at −80°C for future use. Steps hereafter were carried out at 4°C. Membranes containing expressed TRPY1 were prepared by thawing the cells on ice and breaking them using a M110Y microfluidizer (Microfluidics) in homogenization buffer containing 25 mM Tris-HCl, pH 8.0, 300 mM Sucrose and 5 mM EDTA and protease inhibitor cocktail (Sigma). Membranes were then pelleted by first discarding cellular debris with 3,000 x *g* and 14,000 x *g* centrifuge runs, then 100,000 x *g* ultra-centrifugation to pellet the membranes before storing them in 5 mM Tris-HCl, pH 8.0, 300 mM Sucrose and 1 mM PMSF. The membranes were solubilized in 20 mM HEPES pH 8.0, 150 mM NaCl, 10% glycerol, 1% digitonin, 2 mM TCEP, and 1 mM PMSF for 2 h. Insoluble material was removed via ultra-centrifugation at 100,000 x *g* and the solubilized TRPY1 was purified by binding to 1D4 antibody coupled to CnBr-activated Sepharose beads. The beads were used to make a column and extensively washed with Wash Buffer containing 20 mM HEPES pH 7.5, 150 mM NaCl, 0.01% glyco-diosgenin and 2 mM TCEP. The protein was eluted from the column with Wash Buffer supplemented with 3 mg mL^-1^ 1D4 peptide and concentrated to 4.3 mg mL^-1^ using a 100-kDa concentrator (Millipore) before further purifying by size exclusion chromatography using a Superose 6 increase 10/300 GL column (GE Healthcare). The eluted protein was concentrated to 2 mg mL^-1^ and used for vitrification.

### Specimen preparation and cryo-EM data acquisition

Prior to preparing cryo-EM grids, purified protein was incubated with 2 mM CaCl_2_ for 10 min. This sample was blotted once (3.5 mL per blot) onto glow discharged 200 mesh Quantifoil 1.2/1.3 grids (Quantifoil Micro Tools) at 4°C and 100% humidity and plunge frozen in liquid ethane cooled to the temperature of liquid nitrogen (Thermo Fisher Vitrobot Mark IV). Cryo-EM images were collected using a 300 kV Thermo Fisher Krios microscope equipped with a Gatan K3 direct detector camera in super resolution mode. Forty frame movies were collected with a total dose of 45 e/ Å^2^ and a super resolution pixel size of 0.53 Å/pix. Defocus values of the images ranged from −0.5 to −2.5 μm.

### Image processing

Relion 3.1 (version: beta-commit-b86482)^50–52^ was used for data processing of 11,099 super-resolution image stacks (Extended Data Fig. 3). Relion’s implementation of MotionCor2^50,53^ was employed to compensate for beam-induced movement and to set the pixel size to 1.06 Å following 2×2 binning. CTFFIND-4.1^54^ was used to estimate CTF parameters. Laplacian-of-Gaussian autopicking was performed on a small subset (100) of the motion-corrected images followed by two rounds of 2D classification to obtain representative templates for autopicking from the full dataset. Thus, 20,947 initial particles were autopicked, which after two rounds of 2D sorting yielded the best representative template for autopicking on the complete dataset. Autopicking on complete dataset yielded ~3.1 million particles. These particles were 4x-binned and subjected to two rounds of 2D sorting to discard false positives and bad particles. The remaining class averages containing 254,509 particles showed clear signs of being originated from a tetrameric TRP-like channel. These particles were 2x-binned and used to generate an initial model that was subsequently used as a reference to perform a 3D classification with six classes. One class among these six, containing 100,918 particles, aligned well and therefore was better resolved than the rest. Using the map from this class, a loose mask was generated surrounding the tetramer and detergent belt, and thereafter used for 3D refinement and 3D classification of 100,918 unbinned particles applying C4 symmetry and mask. One among the three 3D classes generated, containing 55,593 particles, was subsequently 3D refined to obtain a 3.93 Å map which was then subjected to CTF refinement with corrections for higher-order aberration, anisotropic magnification and per particle defocus, and Bayesian polishing. 3D refinement after Bayesian polishing yielded a 3.35 Å map. A second round of CTF refinement and Bayesian polishing, and a subsequent 3D refinement generated a 3.19 Å map which after postprocessing yielded a 3.08 Å map. This 3.08 Å map was used to build atomic models. The local resolution map was generated using Relion^50^.

### Model building and refinement

An initial model for a single subunit of the full length TRPY1 was generated using the online server ITASSER^55,56^. This model was roughly partitioned into cytosolic, transmembrane and C-terminal domains and rigid body docked into their respective densities. Except for the transmembrane domain, which showed satisfactory fitting, the other two domains did not fit well and therefore were manually adjusted in COOT^57^ to fit the density. The quality of density was comparatively lower in the C-terminal domain and therefore extra care was taken to account for correct registers in the side chain assignments. To assign side chains in the C-terminal helix CH1 and the loop thereafter, spanning Lys517-Asp573, confidence was derived from the distinctive identity of the bulky side chains from Lys538, Arg541, Arg544, Arg545, Tyr548, Arg550, Tyr561, Trp565, Tyr571 and the kinks produced by Pro530 and Pro564. Due to poorer density of CH2, the original I-TASSER model that predicts a helix in this region was fitted. A model for the PI(3)P lipid was built in COOT and thereafter the eLBOW^58^ tool from the PHENIX^59^ software package was used to generate restraints for refinement. Models for Ca^2+^ ions were generated in COOT and were kept unlinked to the protein during refinement. The complete model was then iteratively refined using phenix.real_space_refine from the PHENIX^59^ software package with secondary structure and NCS restraints for the protein and eLBOW-generated restraints for PI(3)P. In the final model, side chains for Leu216, Phe225, Leu572, Asp573 and Lys517-Ser529 were pruned due to insufficient density. Model validation was carried out in Molprobity^60^ and EMRinger^61^. For cross-validation, each final model was randomized by 0.5 Å in PHENIX^59^ and refined against a single half map. These models were converted into volumes in Chimera^62^ and then EMAN2.1^63^ was used to generate FSC curves between these models and each half map as well as between each final model and the final maps. HOLE^64^ was used to generate the pore radii. Electrostatic potential was calculated using APBS-PDB2PQR^65^. Figures were prepared using PyMOL^66^ and Chimera^62^ software.

### TRPY1 molecular dynamics simulations and analyses

The cryo-EM structure of TRPY1 generated in the current study (PDB: 6WHG) was modified as described below and thereafter used for all MD simulations and their associated analyses. The cryo-EM structure has seven missing segments in each chain due to a lack of electron density, from residues Met1 to Asn24 (segment 1; N-terminus), Asp55 to Glu65 (segment 2; loop in cytosolic skirt), Leu216-Phe225 (segment 3; pre-S1 elbow), Pro323-Lys327 (segment 4; loop between S3 and S4 helices), Ala487-Ser529 (segment 5; TRP-CTD linker), Leu572-Ser580 (segment 6; part of C-terminus), and Asp608-Glu675 (segment 7; C-terminal tail). Segments 1 and 7 were not included in our models. Segments 3, 4, and 6 were built using the interface to the Modeler server in Chimera^62^. The top result for each was chosen and was minimized in vacuum for 10,000 steps in NAMD^67,68^ with constraints placed on all but the modeled segments (this and subsequent harmonic constraints used *k* = 1 kcal mol^-1^ Å^-2^). Real space refinement was performed on the resulting model to better fit the cryo-EM map. Segment 2 was built in COOT^57^, based on the cryo-EM map with C4 symmetry applied and no sharpening. This was then minimized in vacuum for 50,000 steps in NAMD with all but segment 2 constrained. Because segment 5 was predicted to contain a helix based on its sequence, this segment was submitted to the Phyre2 homology-modeling server^69^. The top model from Phyre2 contained an amphipathic helix and was inserted into the TRPY1 structure using COOT. This segment was then minimized in vacuum for 10,000 steps in NAMD with all but segment 5 constrained. This was done for a single chain. To insert the missing segments for the remaining three chains, the completed chain was aligned to the other three chains in COOT, and the coordinates of the missing segments were saved. These were then inserted into the remaining chains, and all completed chains were saved as a single model, which was then vacuum minimized for 50,000 steps in NAMD, with all but the missing segments constrained.

The TRPY1 structure (PDB: 6WHG) also contains four PI(3)P molecules bound as ligands with a partial missing tail. To complete these ligands, the internal coordinates from the SAPI13 molecule in the toppar_all36_lipid_inositol.str CHARMM parameter file were used with the autopsf plugin. The entire structure was then vacuum minimized in NAMD for 50,000 steps with all but the PI(3)P molecules constrained. The final coordinates were saved, and this entire structure was embedded in a 150 × 150 Å membrane patch that contained 50% POPC, 18% POPA, 16% POPS, and 16% POPI. The membrane patch was generated using the Membrane Builder function on the CHARMM-GUI server. The transmembrane region was placed based on hydrophobic regions of the core helices, proximity of tryptophan residues to lipid head groups, and previous simulations performed on TRP channels^70,71^. Then, the solvate plugin in VMD^72^ was used to add TIP3P water molecules to the system, and the autoionize plugin was used to neutralize the system and add KCl to a final concentration of 150 mM.

Simulations were ran using NAMD 2.12^67,68^ and the CHARMM36 force field^73^. The TIP3P^74^ explicit model of water was used. Long-range electrostatic forces were computed using the Particle Mesh Ewald method with a grid point density of >1 Å^-3^, and a cutoff of 12 Å was used for van der Waals interactions. The SHAKE algorithm was used with a timestep of 2 fs. The *NpT* ensemble was used at 1 atm with a hybrid Nosé-Hoover Langevin piston method and a 200 fs decay period, with a 50 fs damping time constant. The completed simulation system was minimized for 1,000 steps, followed by a lipid-melting step in which all but the lipid tails were constrained for 0.5 ns of equilibration. This was followed by 0.5 ns of equilibration in which the protein was constrained and the water, lipids, and ions were free. Next, constraints were released on the modeled segments of the protein for 10 ns, allowing them to equilibrate with their surroundings. Finally, all constraints were lifted and the system was ran for 20 ns using a Langevin damping coefficient of γ = 1, after which γ was set to 0.1 for 100 ns of free equilibration. All analyses were performed on the final 100 ns for every simulation.

A total of 5 simulations were prepared and ran as described above. The first four are variations of a “vertical” configuration, in which the CTD-linker was modeled with its helix segment vertically across the membrane (Extended Data Fig. 10). The first is the All-Bound system, in which all Ca^2+^ and PI(3)P sites are occupied. The second is the None-Bound system, in which all Ca^2+^ and PI(3)P are removed. For the third system, labeled No-Inh, only the non-inhibitory (cytosolic) Ca^2+^-binding sites remain occupied, while the PI(3)P and inhibitory (luminal) Ca^2+^ ions were removed. Finally, in the fourth system, a gain of function Y473H mutation was introduced to the non-inhibitory system using VMD’s Mutate Residue plugin.

An additional system was built based on the “horizontal” configuration, in which the amphipathic helix of segment 5 (TRP-CTD linker) was modeled as laying horizontally at the lipid-cytosolic interface (Extended Data Fig. 10). This system, besides segment 5, is identical to the All-Bound system of the vertical configuration.

The network analysis was performed following standard protocols^35^. First, a dynamic network is created from the MD trajectory in VMD using Carma^75^, a program that calculates correlation between atoms in an MD simulation. Optimal and suboptimal paths are calculated by first choosing a source and sink node, and the communication pathways between these nodes are calculated for the minimal number of nodes. Those nodes that occur in the most sub-optimal paths are critical nodes for allosteric communication between source and sink nodes. All visualization of molecular images and allosteric pathways was performed in VMD. Ion and water densities were calculated using the VolMap tool plugin in VMD. The average density was computed and combined for all frames with no weight after alignment. These data, given in number of atoms/Å^3^, were then converted to a contour map and reported in molarity.

For each frame of the given simulation, RMSD was calculated based on the alignment of a selection of Cα atoms to the same selection of atoms in the first frame of the trajectory. In this way, the RMSD analysis of every system was done based on the same starting structure.

The average pore radius (Fig. 5) was calculated using the program HOLE^64^. After alignment of the protein to the first frame of the trajectory, HOLE calculates the pore radius in every frame of the trajectory along the path of the pore in the *z*-dimension. Next, these values are averaged throughout the trajectory at each position in the pore, with error bars in Fig. 5 indicating the standard deviation.

## Data availability

The cryo-EM density map and the atomic coordinate of PI(3)P and Ca^2+^-bound full-length TRPY1 in detergent are deposited into the Electron Microscopy Data Bank and Protein Data Bank under accession codes EMD-21672 and 6WHG. To avoid repetition, we have not submitted the apo condition map and model in the databases.

## Acknowledgements

We thank Sabine Baxter for assistance with hybridoma and cell culture at the University of Pennsylvania Perelman School of Medicine Cell Center Services Facility. We acknowledge the use of instruments at the Electron Microscopy Resource Lab and at the Beckman Center for Cryo Electron Microscopy at the University of Pennsylvania Perelman School of Medicine. We also thank Darrah Johnson-McDaniel for assistance with Krios microscope operation. This work was supported by grants from the National Institute of Health (R01GM103899 and R01GM129357 to VYM-B). Use of TACC-Stampede2 and PSC-Bridges supercomputers was supported by the National Science Foundation through XSEDE (XRAC MCB140226 to MS). Use of resources at the Ohio Supercomputer Center was supported by grants PAS1037 and PAA0217 to MS. CN was supported by an OSU/NIH molecular biophysics training grant (T32GM118291).

---

**Extended Data Figure 1:**
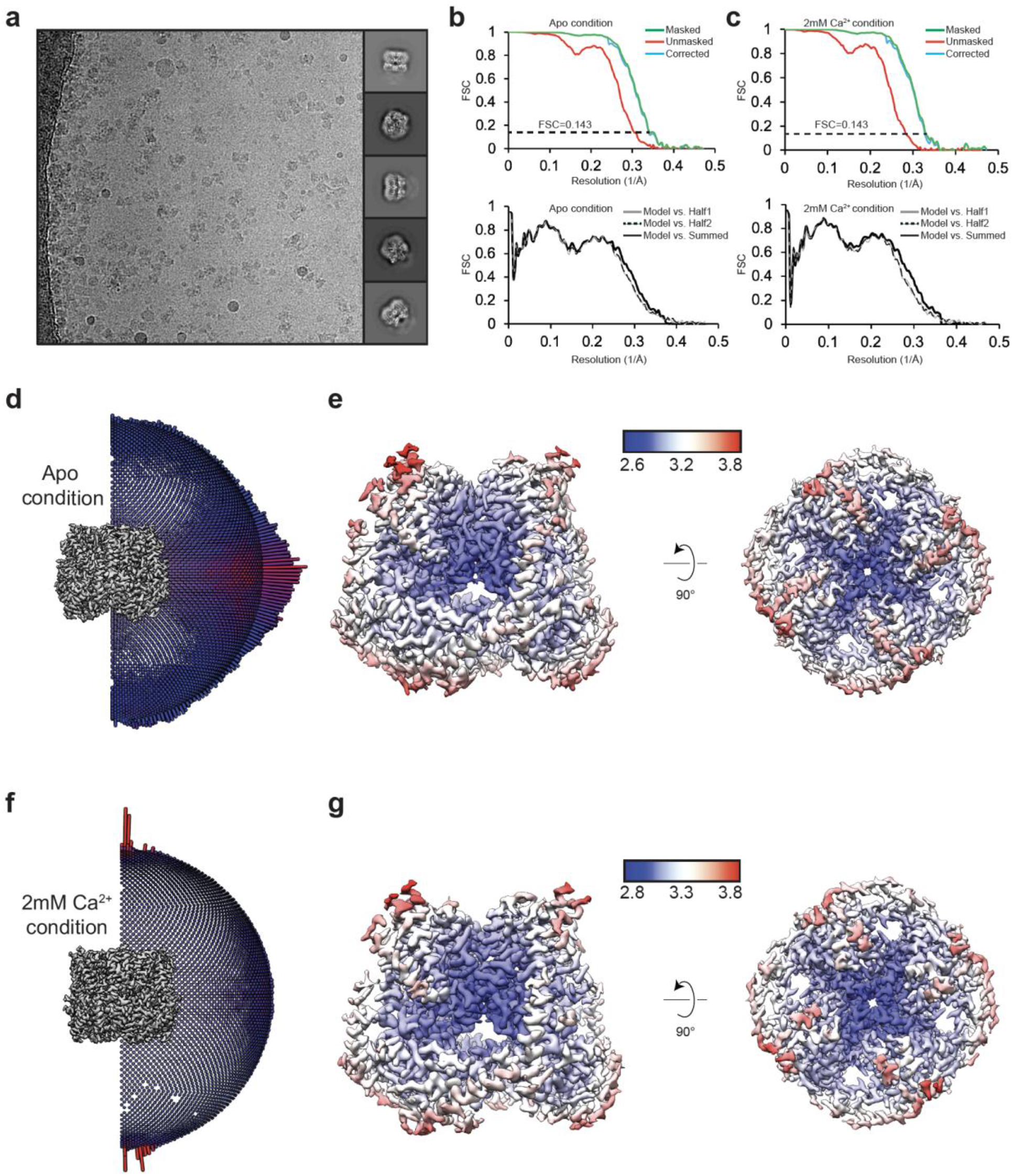
EM summary of TRPY1 in detergent. **a,** Representative cryo-EM micrograph collected at 2.8 μm defocus (left) and 2D class averages generated from cryo-EM data (right). **b, c,** Map FSC and model validation curves for TRPY1 in **(b)** apo and **(c)** 2mM Ca^2+^ conditions. **d, f,** Angular distribution of the particles used for 3D reconstruction for the final map of TRPY1 apo **(d)** and 2mM Ca^2+^ conditions **(f)**. **e, g,** Preferred orientation of the particles is represented by either a higher or lower number as denoted by red or blue rods, respectively. Maps from TRPY1 apo condition at Chimera^62^ threshold 0.03 **(e)** and 2mM Ca^2+^ condition at threshold 0.034 **(g)** shown in side (left) and cytosolic (right) views and colored according to the local resolution calculated by RELION^52^. The bar displaying blue, white and red colors denotes resolution measured in Å.

**Extended Data Figure 2:**
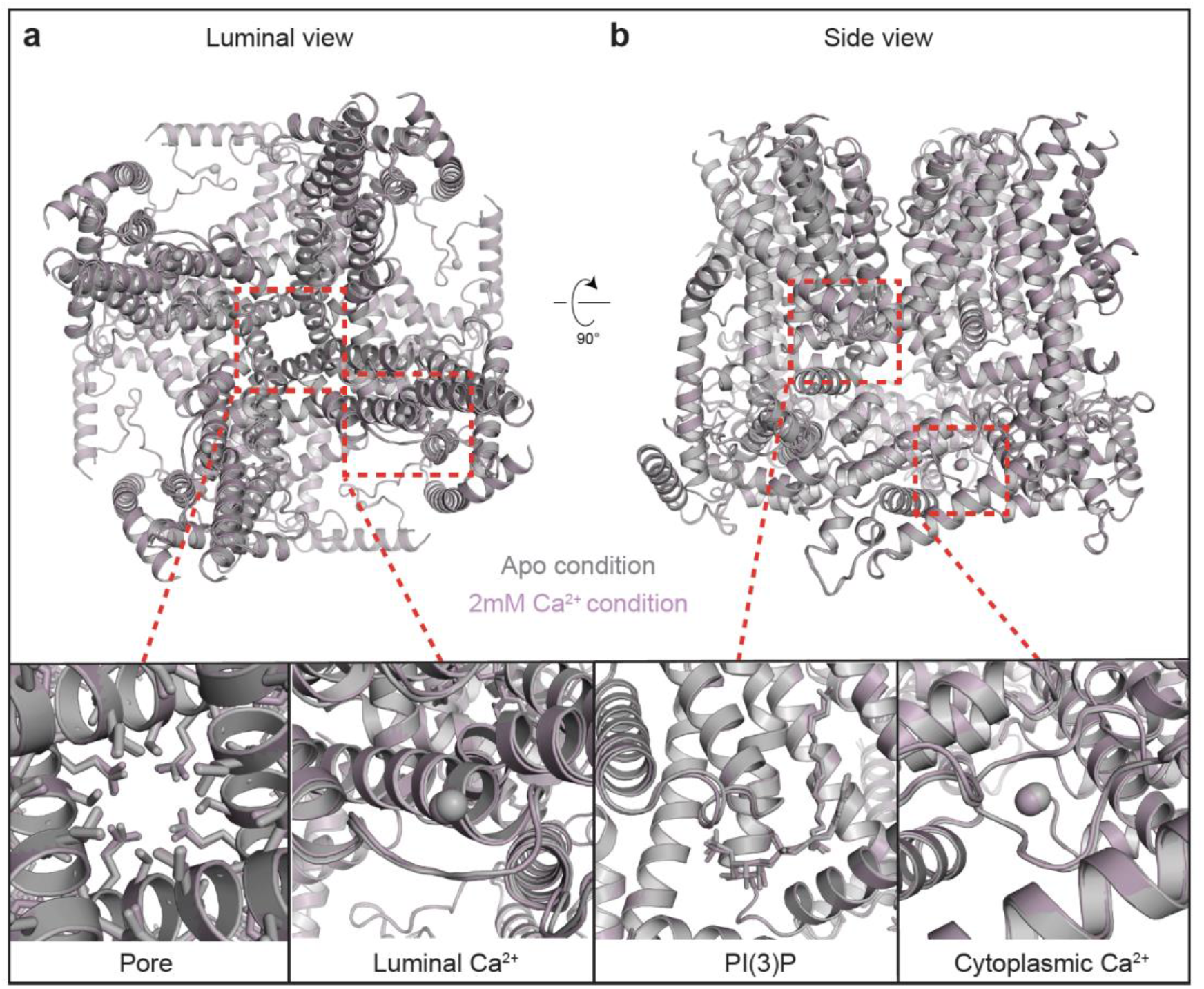
Comparison of TRPY1 structures in apo and 2mM Ca^2+^ conditions. **a, b,** Models for **(a)** TRPY1 apo (grey) and **(b)** 2mM Ca^2+^ conditions (plum) are overlaid and shown in the vacuolar lumen and side views (top). The regions encompassing pore, luminal Ca^2+^, PI(3)P and cytosolic Ca^2+^ are zoomed in to show structural similarity among the two models.

**Extended Data Figure 3:**
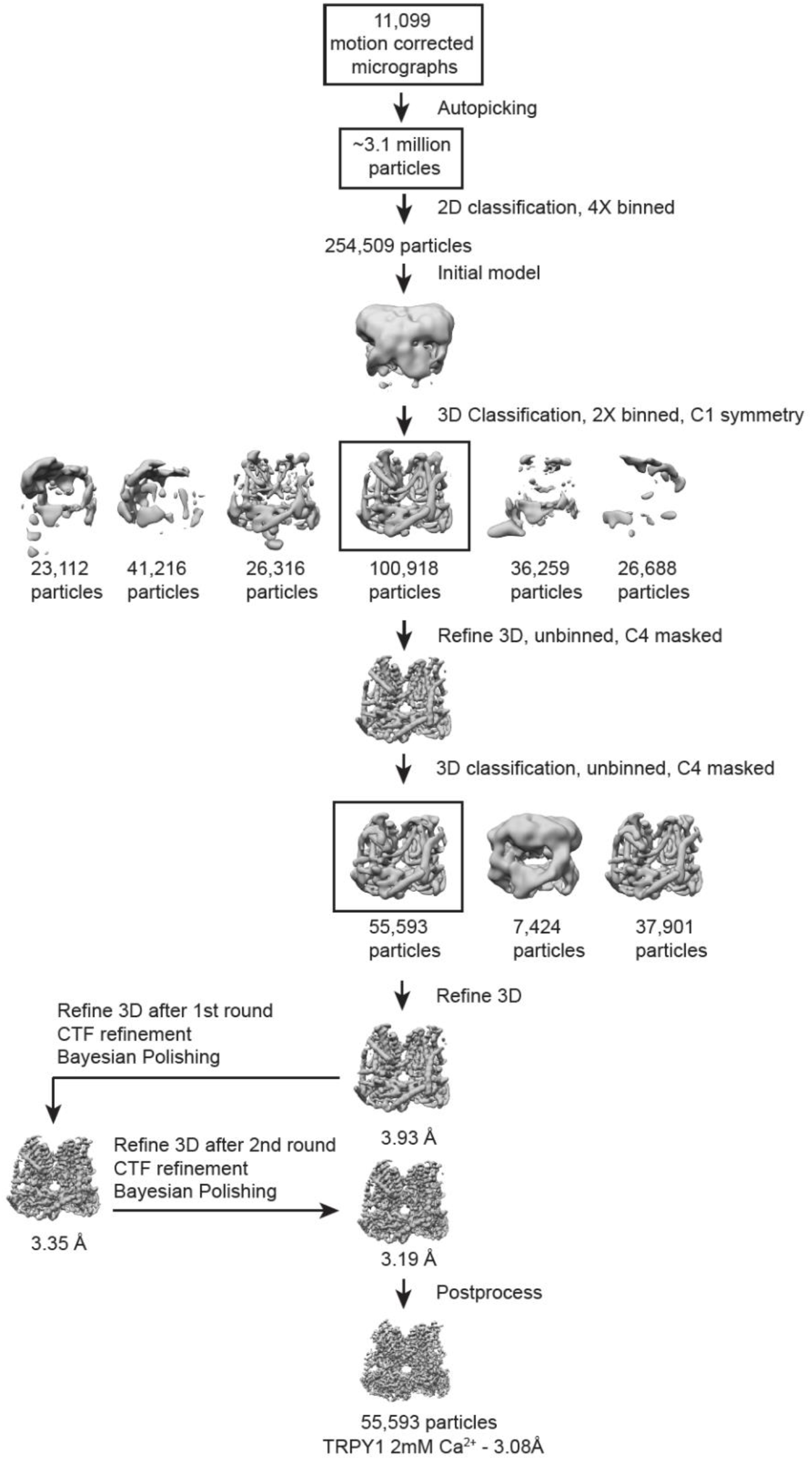
Data processing workflow resulting in the final TRPY1 map in 2mM Ca^2+^ condition used for model building.

**Extended Data Figure 4:**
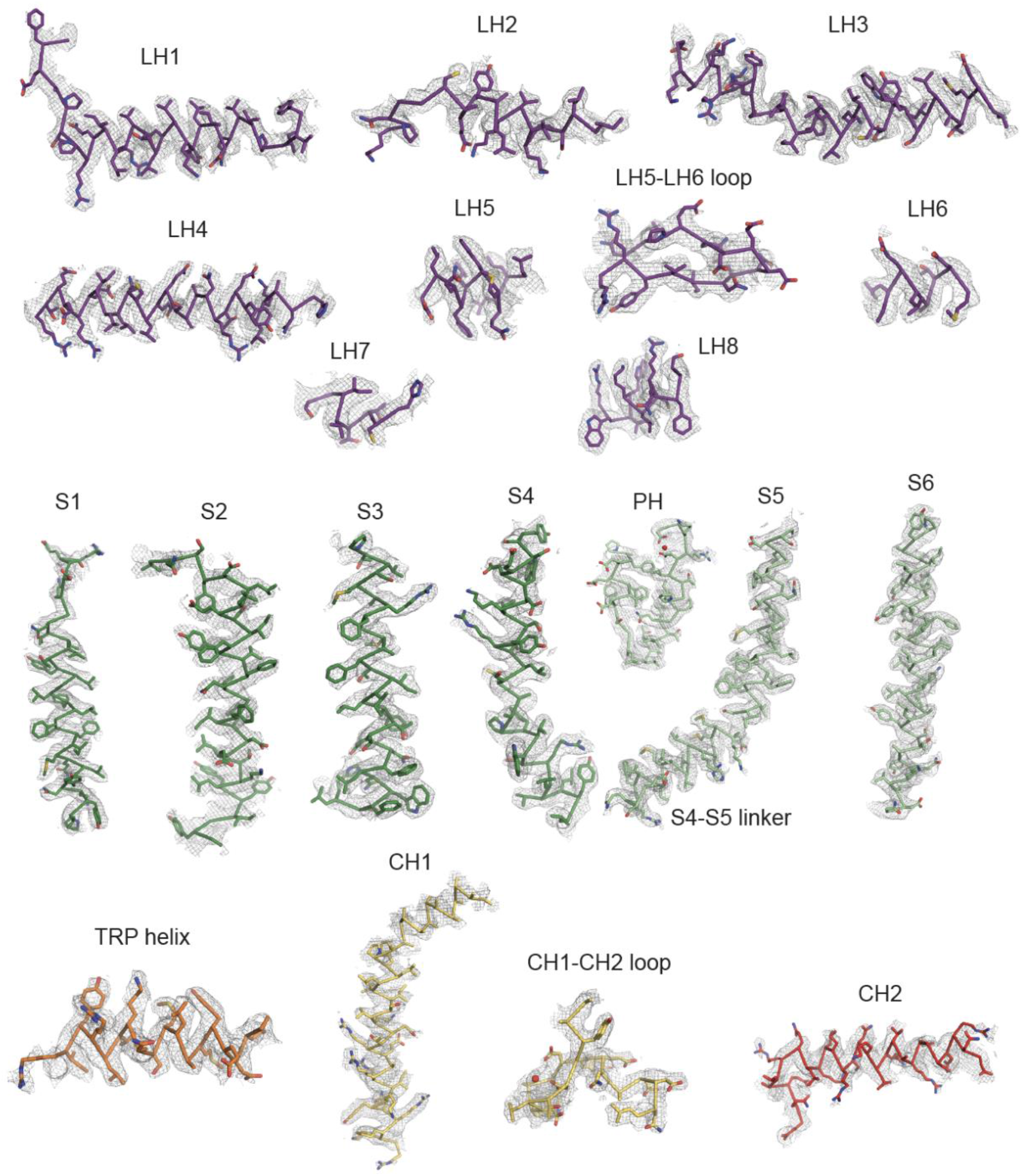
Representative densities from the TRPY1 map in the presence of 2mM Ca^2+^. Densities are contoured at sigma level 1.25.

**Extended Data Figure 5:**
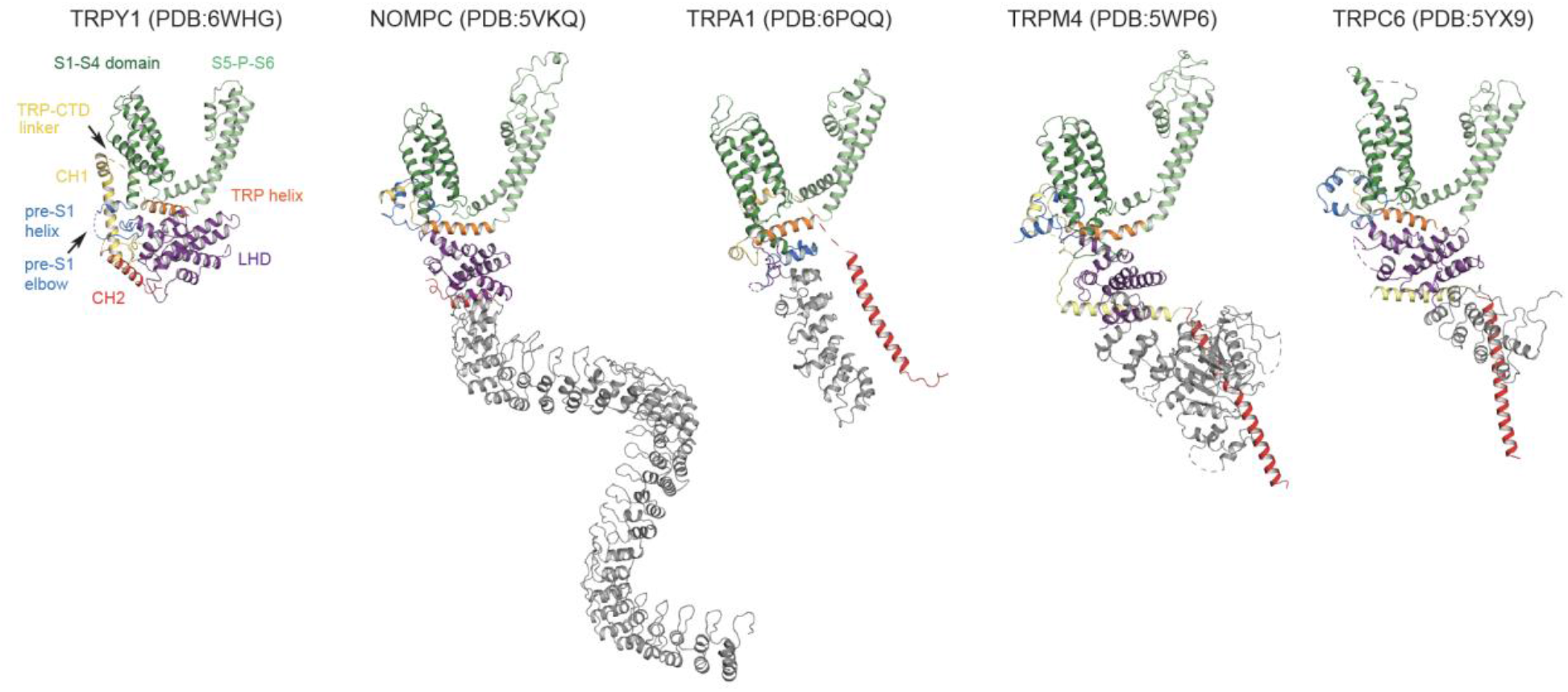
Comparison of TRPY1 structure with other TRP channels. A subunit from the TRPY1 structure derived from the current study (extreme left) with domains colored differently and labelled. Subunits from the TRP channels possessing structural similarity with TRPY1 are displayed with their domains color-matched with TRPY1. The extra domains which are absent from the TRPY1 structure are colored in grey.

**Extended Data Figure 6:**
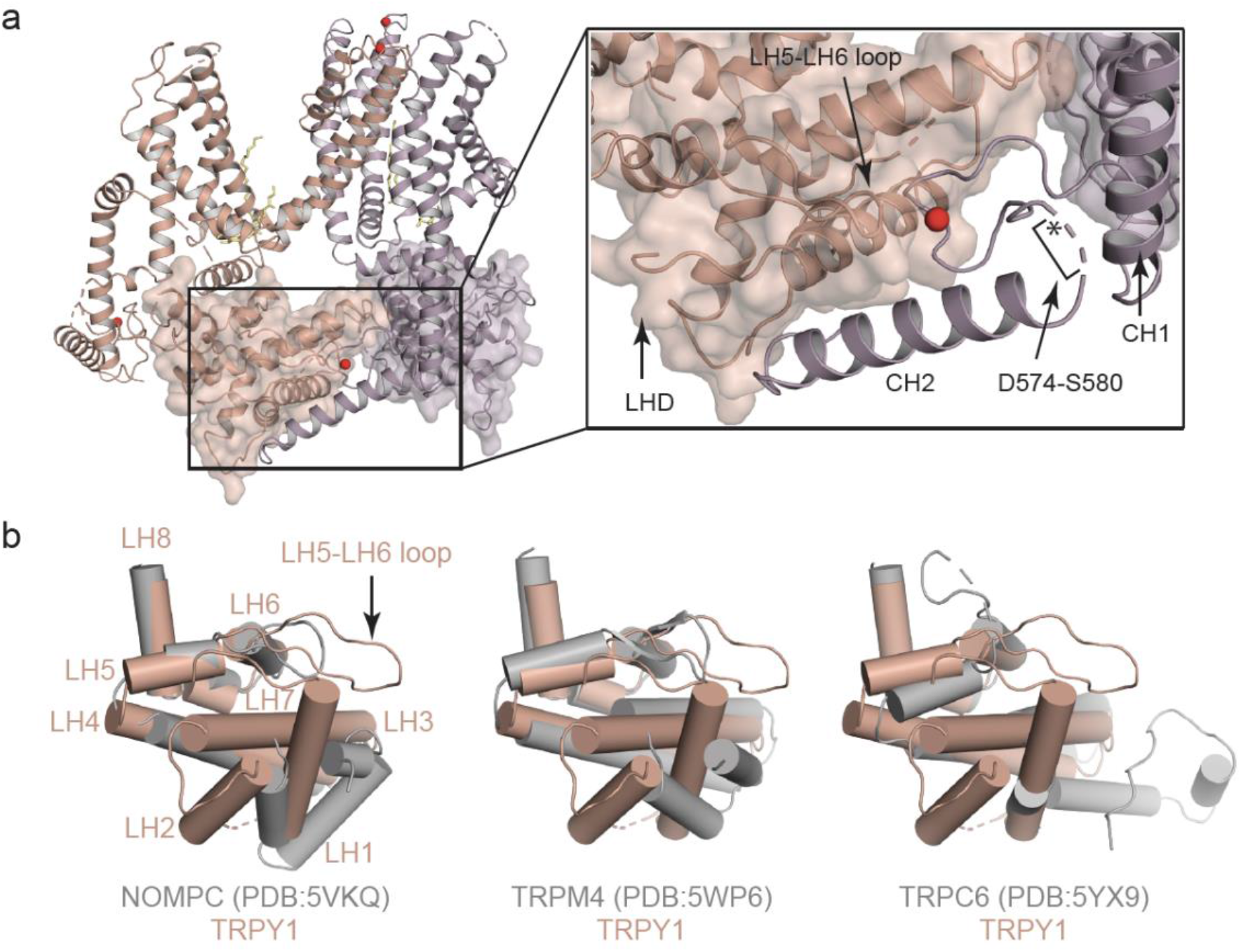
Distinct structural features of TRPY1 LHD. **(a)** Two subunits (salmon and plum cartoons) of TRPY1 shown to display close interaction of LHD and CTD from neighboring subunits. Both LHDs are also represented as transparent surface generated from the model. The cytosolic Ca^2+^ is in red. The position of C-terminal acidic patch is marked by an asterisk. Due to possible flexibility, density between D574-S580 is absent and not modeled. **(b)** TRPY1 LHD structure in comparison to LHDs from NOMPC, TRPM4 and TRPC6.

**Extended Data Figure 7:**
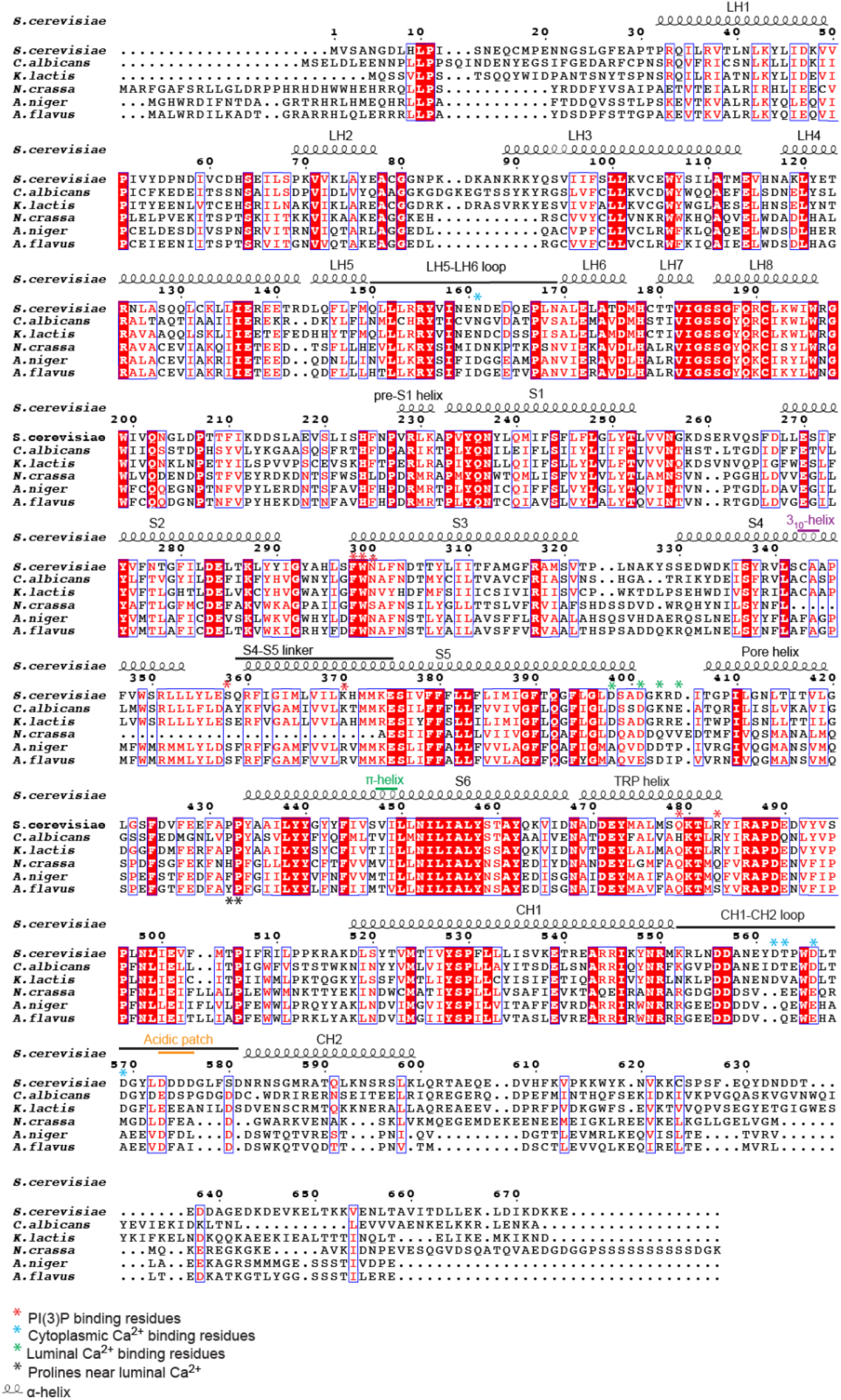
Sequence alignment of *S. cerevisiae* TRPY1 with other fungal homologs. The secondary structure annotation related to helices and loops are generated from the *S. cerevisiae* TRPY1 model built in the current study. Sequence numbering is based on TRPY1. Asterisks in different colors denote important regulatory residues as revealed in our TRPY1 structure.

**Extended Data Figure 8:**
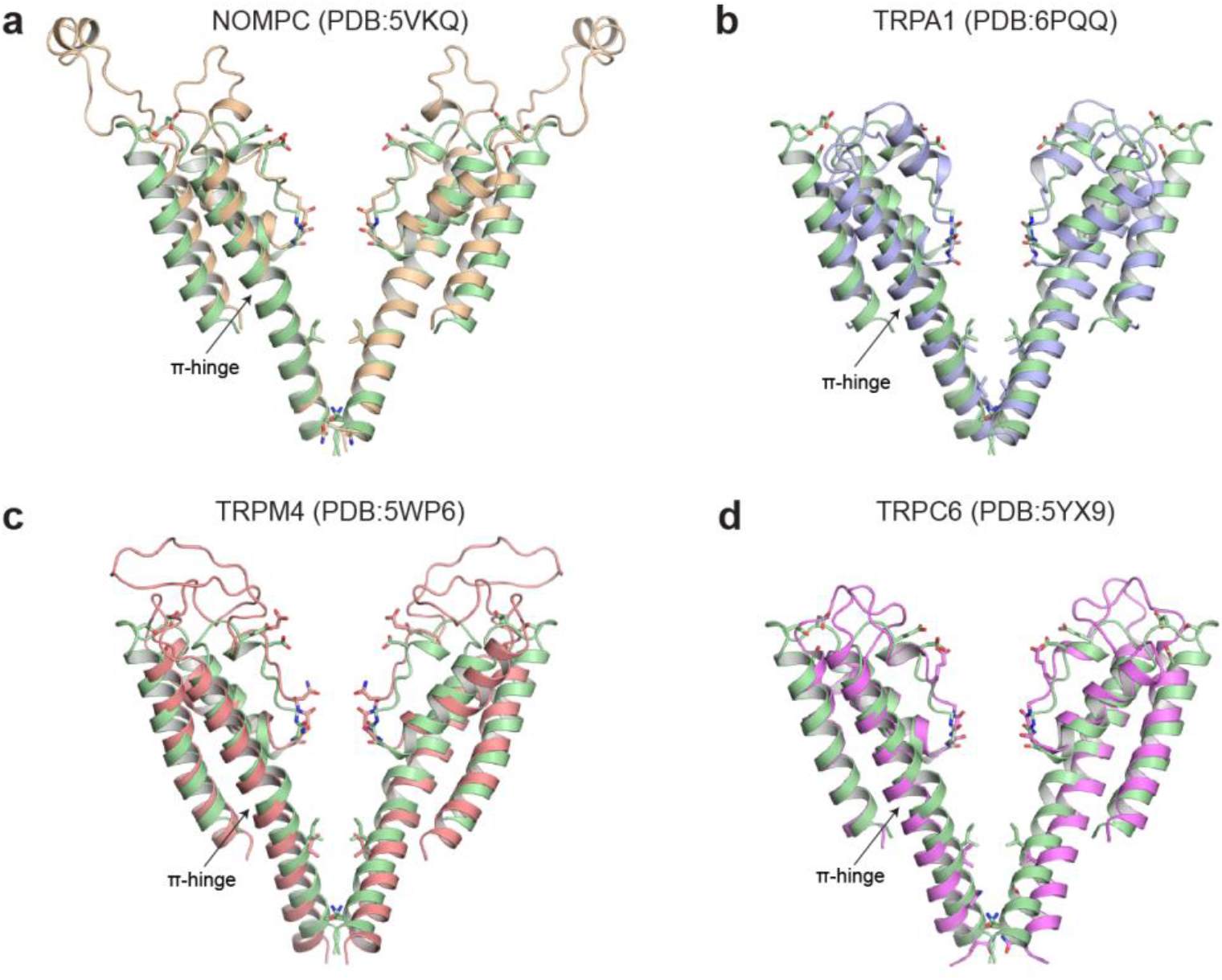
Comparison of TRPY1 ion permeation pathway with other TRP channels. Drosophila NOMPC and TRPA1, TRPM4 and TRPC6 from human are pore-aligned on S5 and S6 helices to TRPY1 in order to show similarity. Models are colored as: TRPY1: pale green, NOMPC: wheat, TRPA1: light blue, TRPM4: salmon and TRPC6: violet.

**Extended Data Figure 9:**
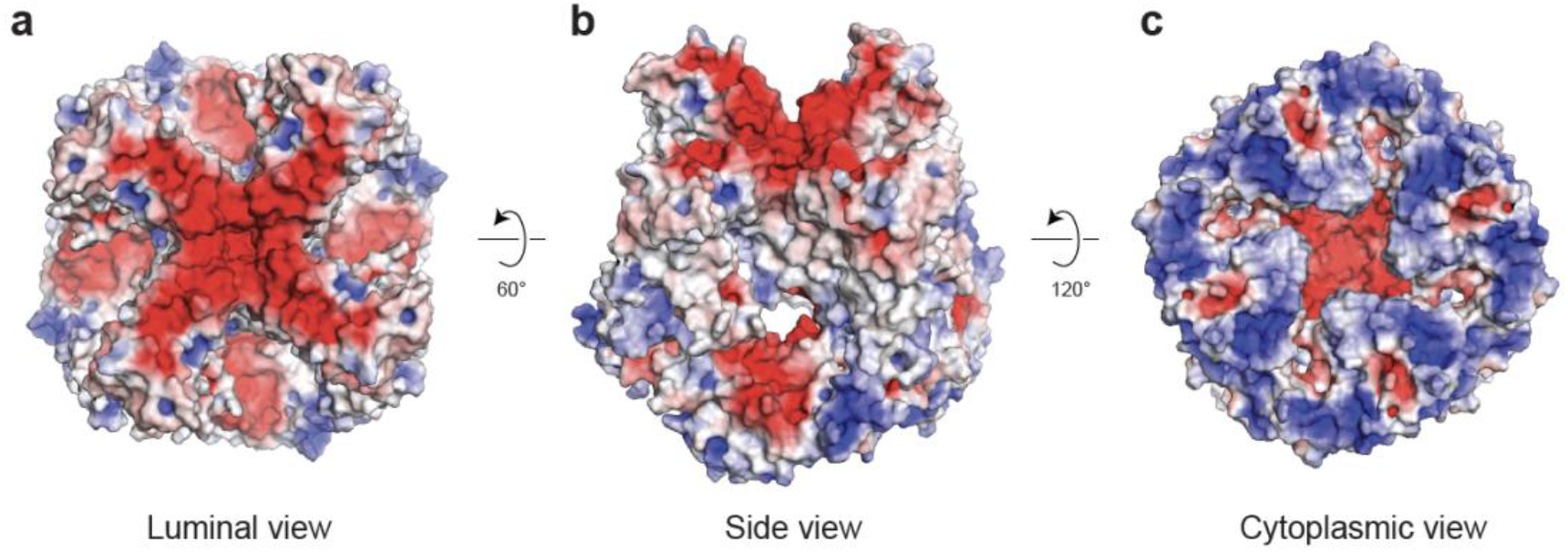
Electrostatic potential surface. Luminal **(a)**, tilted side **(b)** and cytosolic **(c)** views are shown for the electrostatic potential surface generated from TRPY1 2mM Ca^2+^ condition model. Negative and positive electrostatic potentials are colored as red and blue, respectively. Ca^2+^ ions and PI(3)P lipids were excluded before calculating the surface.

**Extended Data Figure 10:**
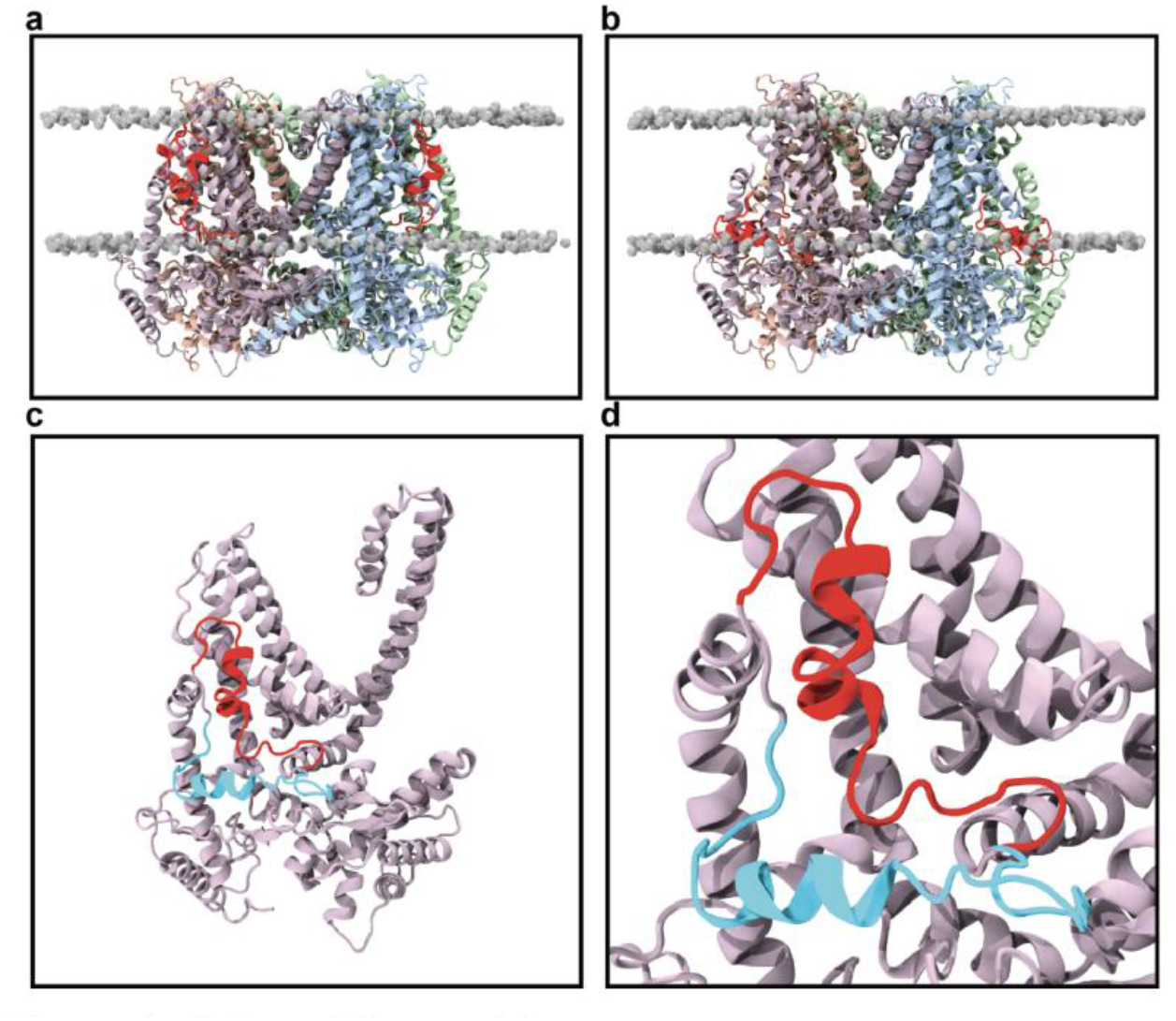
Comparison of the TRP-CTD linker in the vertical and the horizontal configurations. The vertical configuration **(a)** and the horizontal configuration **(b)** with the TRP-CTD linker (residues Ala487-Ala516) shown in red, and phosphorous atoms of the lipid bilayer shown in grey spheres. **c-d,** Chain A TRP-CTD linker shown in red for the vertical configuration, and in cyan for the horizontal configuration.

**Extended Data Figure 11:**
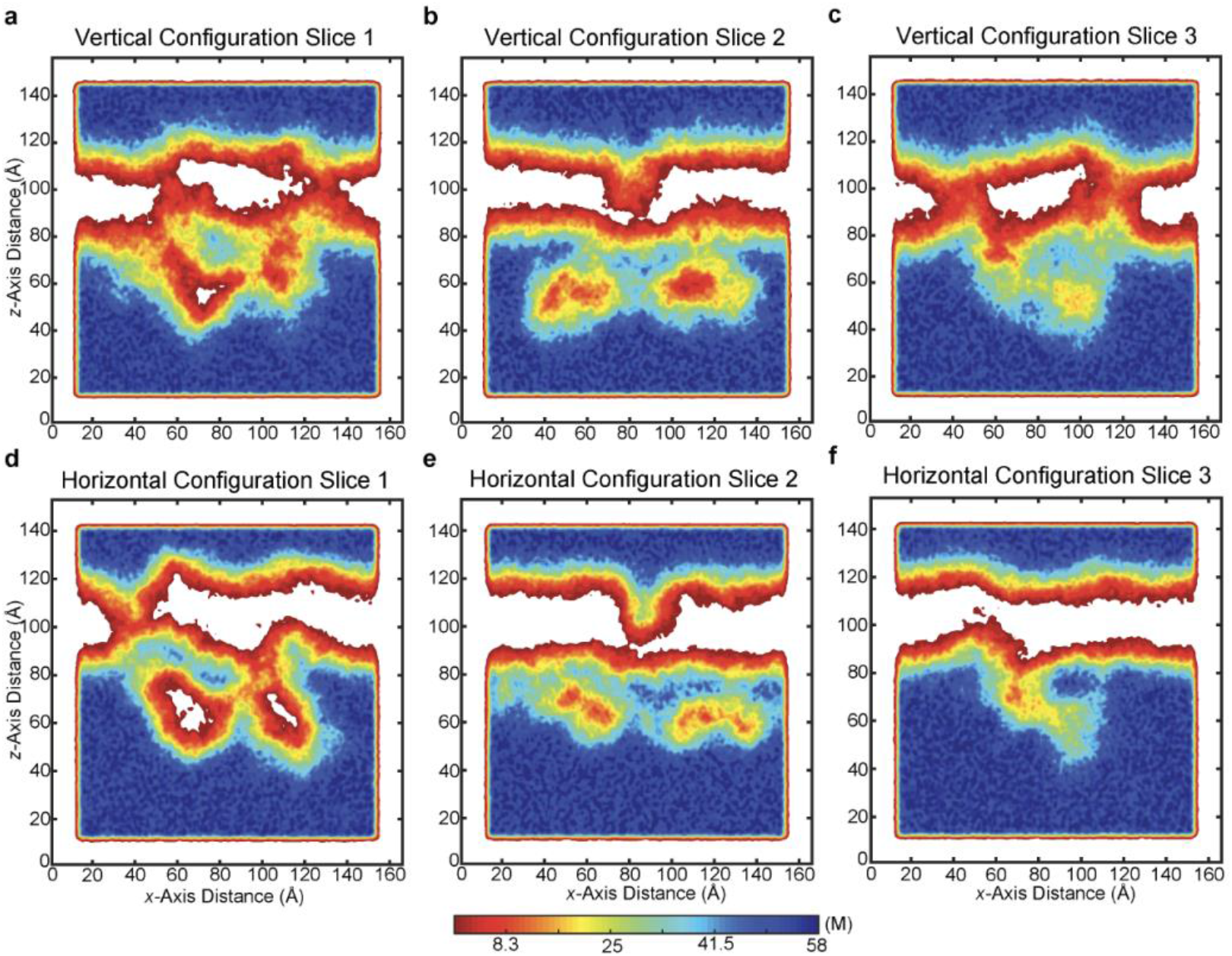
Reduced water leakage through lipid bilayers in the horizontal configuration. **(a-c),** The water density is shown in the *x-z* plane at three different points along the *y*-axis for the vertical configuration. In **(a)** and **(c)**, some water can be seen within the lipid bilayer, indicating that this configuration results in minor leaking across the membrane. **(d-f)**, Water density in the *x-z* plane at the same points along the *y-*axis as in **(a-c)**, but for the horizontal configuration. This configuration results in reduced leaking of water across the lipid bilayer, as seen in **(d)** and **(f)**.

**Extended Data Figure 12:**
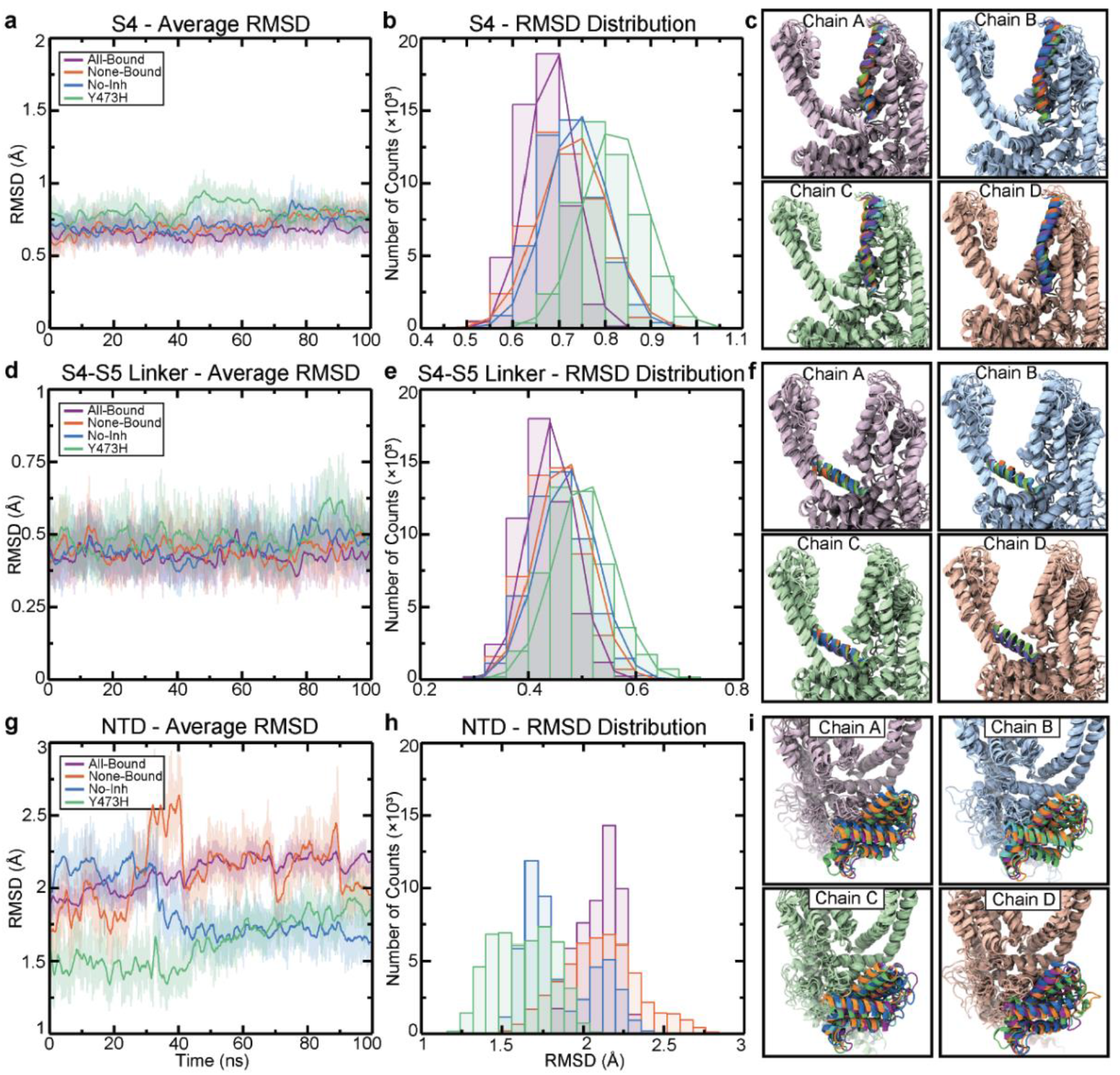
RMSD analysis of the vertical configuration shows differing dynamics between systems with varying Ca^2+^ and PI(3)P bound states. **a,** The average RMSD of the S4 helix for all chains is shown for all systems during the 100 ns trajectory. **b,** Count distributions from (**a**) with associated fits to a normal distribution. **c,** All systems have been aligned based on all of the Cα atoms of the indicated chain other than the S4 helix. Colors correspond to those in (**a**) and (**b**). **d,** The average RMSD of the S4-S5 linker for all chains is shown for all systems during the 100 ns trajectory. **e,** Count distributions from (**d**) as in (**b**). **f,** All systems have been aligned based on all of the Cα atoms of the indicated chain other than the S4-S5 helix. Colors correspond to those in (**d**) and (**e**). **g,** The average RMSD of the N-terminal domain (NTD) helices (residues Pro32-Asp143) for all chains is shown for all systems during the 100 ns trajectories. **h,** Count distributions are shown from (**g**). Fits to a normal distribution were not performed due to irregular distributions and multiple populations exhibited by multiple systems. **i,** All systems have been aligned based on all of the Cα atoms of the indicated chain other than residues Pro32-Asp143. Colors correspond to those in (**g**) and (**h**). All RMSD alignments were performed based on the initial structure, prior to any simulation (shown in cyan in each panel), and the final frame of the trajectory for each system.

**Extended Data Figure 13:**
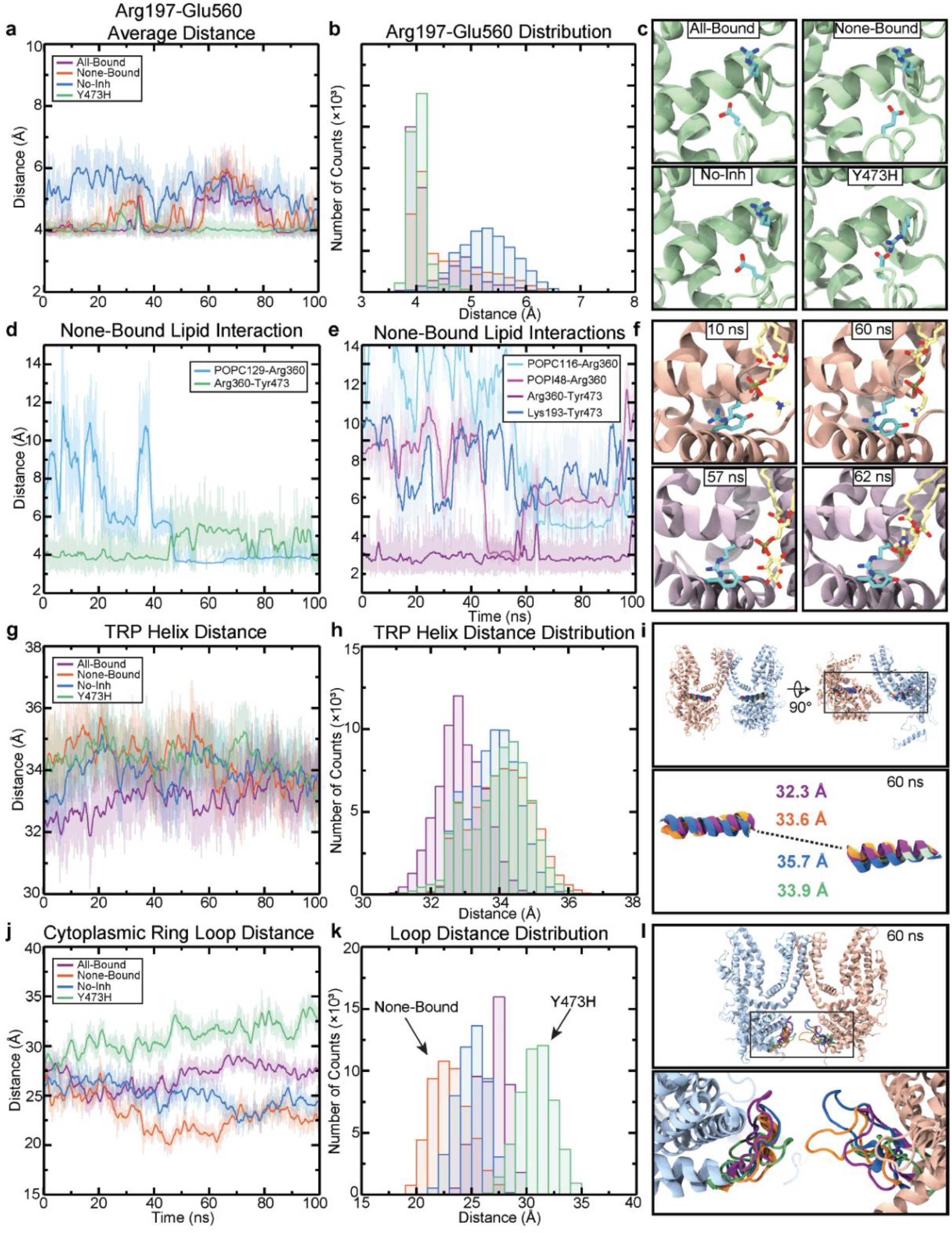
Distance analyses of vertical configuration simulations reveal greater destabilization of key interactions and altered dynamics when PI(3)P or Ca^2+^ is removed. **a,** The average distance of the Arg197-Glu560 interaction for all chains is shown for all systems during 100 ns-long trajectories. **b,** Count distributions from (**a**). **c,** Residues Arg197 and Glu560 are shown for each system at 60 ns. **d,** Distance between Arg360 and Tyr473 in chain D as well as between POPC129 and Arg360 for the None-Bound system. As the POPC lipid approaches Arg360, the Arg360-Tyr473 interaction is destabilized. **e,** In chain A, The Arg360-Tyr473 interaction is destabilized by POPC and POPI lipids, as well as nearby Lys193. **f,** Interactions from (**d**), (top row), and (**e**), (bottom row), are shown at the indicated timepoints. **g,** The distance between the Cα atom of residue Ala469 (at the apex of the TRP helix) of opposing TRP helices is shown. **h,** Count distributions from (**g**). **i,** All systems have been aligned based on all of the Cα atoms of chain B and D other than the TRP helix. Colors correspond to those in (**g**) and (**h**). **j,** The distance between the center of mass of residues Val50-Pro69, comprising the loop that forms the neck of the cytosolic skirt, is shown for chains B and D. **k,** Count distributions from (**j**). **i,** All systems have been aligned based on all of the Cα atoms of chain B and D other than residues Val50-Pro69. Colors correspond to those in (**j**) and (**k**).

**Extended Data Figure 14:**
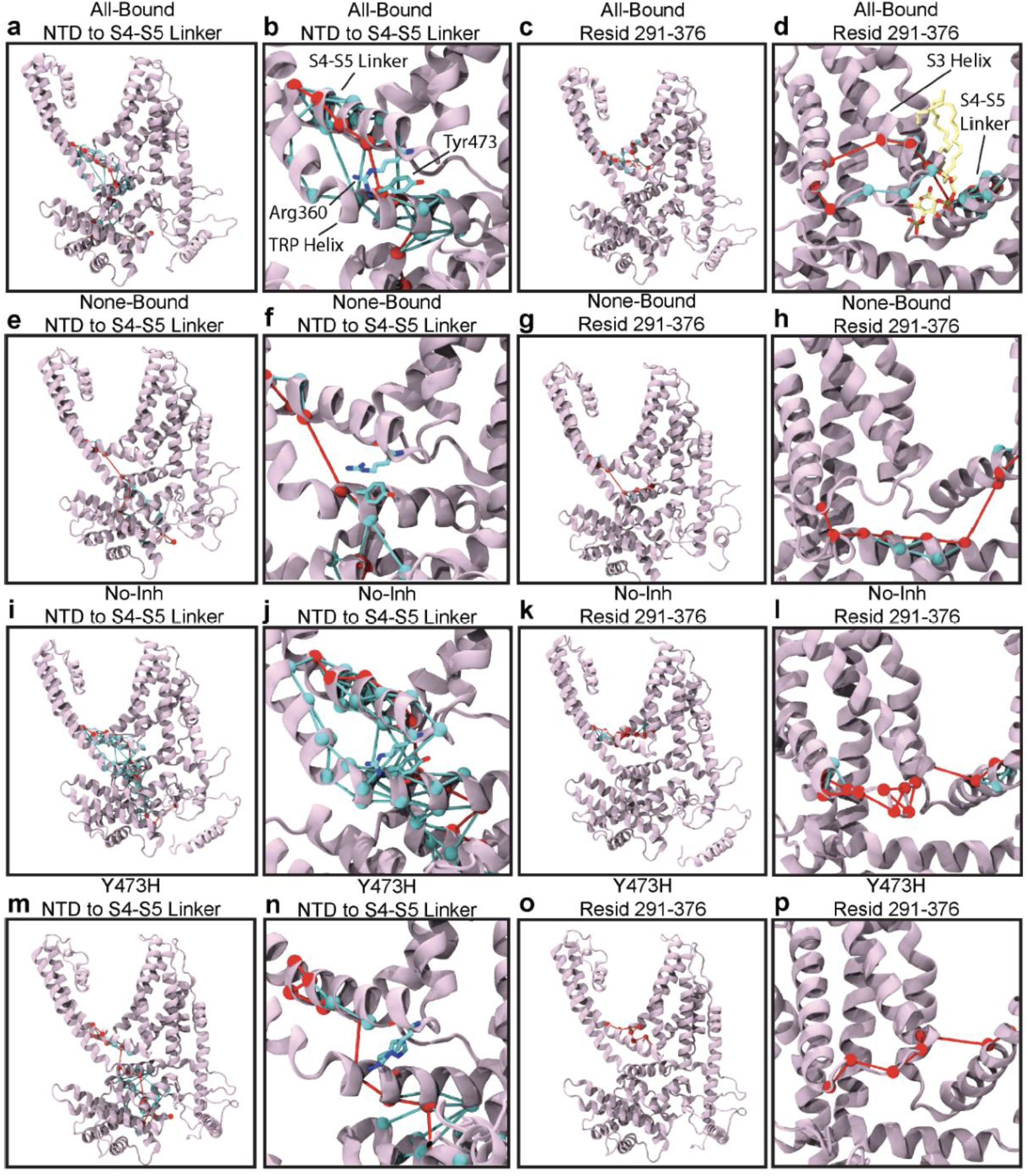
Dynamic network analysis of the vertical configuration simulations show removal of Ca^2+^ and PI(3)P disrupts allosteric communication in TRPY1. All-Bound (**a-b**), None-Bound (**e-f**), No-Inh **(i-j**), and Y473H (**m-n**) optimal (red) and sub-optimal paths (cyan) from the N-terminus to the S4-S5 linker. The optimal path passes through residue Tyr473 only in the All-Bound system, while in the None-Bound and the Y473H systems there is far less communication between the TRP helix and the S4-S5 linker. All-Bound (**c-d**), None-Bound (**g-h**), No-Inh **(k-l**), and Y473H (**o-p**) optimal (red) and sub-optimal paths (cyan) from the S4-S5 helix to residue Ile291 at the base of the S2 helix. When PI(3)P is bound, as in (**c-d**), the optimal path crosses the midpoint of the S2-S3 helices. In all other cases, the optimal path travels through the loop connecting the S2 and S3 helices, or the TRP helix, demonstrating greater communication between the transmembrane helices when PI(3)P is present.

**Extended Data Figure 15:**
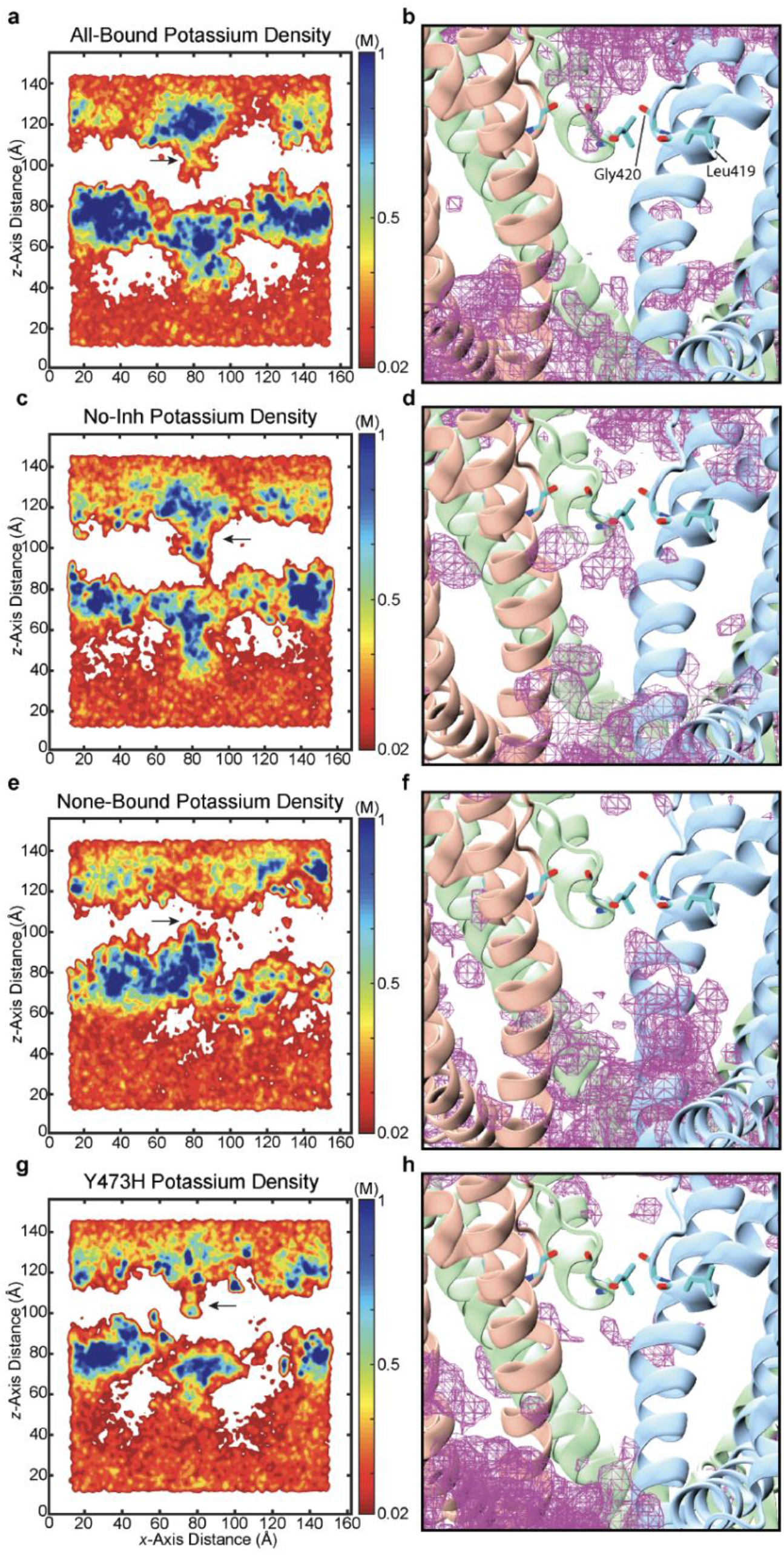
K^+^ density of vertical configuration simulations show that removal of inhibitory Ca^2+^ and PI(3)P increases ion access to the pore. **(a, c, e, g),** The density of K^+^ ions is shown at the origin of the simulation system, corresponding to the center of the pore, for each system in the *x-z* plane. (**b, d, f, h**), K^+^ density represented as a magenta mesh. The Y473H (**h**) and No-Inh (**d**) systems exhibit density past the selectivity filter, shown in licorice representation, while the All-Bound system (**b**) does not. Black arrows indicate the location of residue Gly420.

**Extended Data Table 1:**
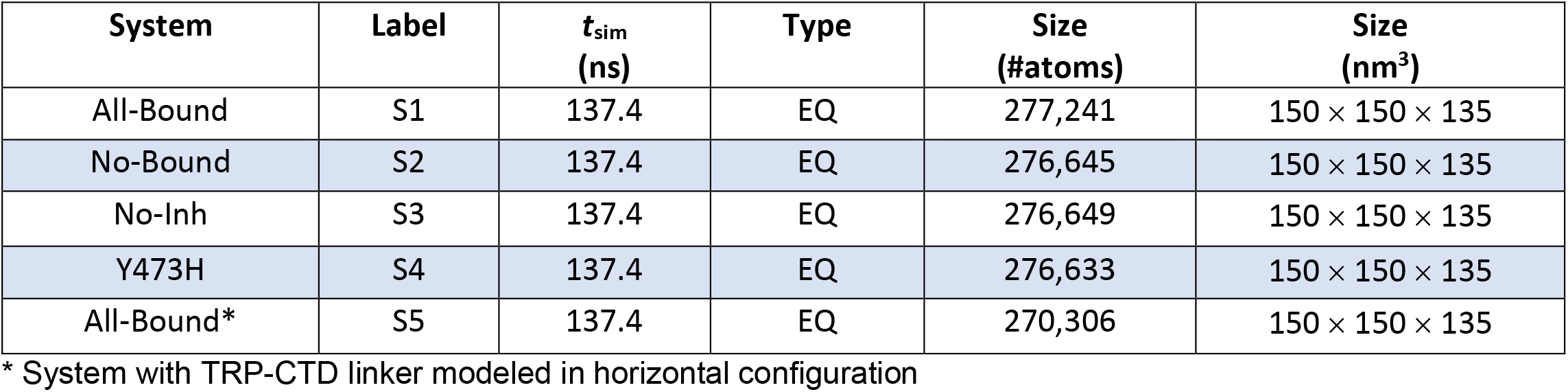
TRPY1 Molecular Dynamics Simulations

## REFERENCES

1. Hohmann S, Krantz M, Nordlander B. Yeast Osmoregulation. In: Häussinger D, Sies HBT-M in E, eds. Methods in Enzymology. Vol 428. Academic Press; 2007:29–45. doi:10.1016/S0076-6879(07)28002-4

2. Hohmann S. Osmotic Stress Signaling and Osmoadaptation in Yeasts. Microbiol Mol Biol Rev. 2002;66(2):300–372. doi:10.1128/mmbr.66.2.300-372.2002

3. Cui J, Kaandorp JA, Sloot PMA, Lloyd CM, Filatov M V. Calcium homeostasis and signaling in yeast cells and cardiac myocytes. FEMS Yeast Res. 2009;9(8):1137–1147. doi:10.1111/j.1567-1364.2009.00552.x

4. Halachmi D, Eilam Y. Cytosolic and vacuolar Ca2+ concentrations in yeast cells measured with the Ca2+-sensitive fluorescence dye indo-1. FEBS Lett. 1989;256(1-2):55–61. doi:10.1016/0014-5793(89)81717-X

5. Chang Y, Schlenstedt G, Flockerzi V, Beck A. Properties of the intracellular transient receptor potential (TRP) channel in yeast, Yvc1. FEBS Lett. 2010;584(10):2028–2032. doi:10.1016/j.febslet.2009.12.035

6. Hellmich UA, Gaudet R. Structural Biology of TRP Channels BT-Mammalian Transient Receptor Potential (TRP) Cation Channels: Volume II. In: Nilius B, Flockerzi V, eds. Cham: Springer International Publishing; 2014:963–990. doi:10.1007/978-3-319-05161-1_10

7. Julius D. TRP Channels and Pain. Annu Rev Cell Dev Biol. 2013;29(1):355–384. doi:10.1146/annurev-cellbio-101011-155833

8. Ramsey IS, Delling M, Clapham DE. AN INTRODUCTION TO TRP CHANNELS. Annu Rev Physiol. 2006;68(1):619–647. doi:10.1146/annurev.physiol.68.040204.100431

9. Rosasco MG GS. TRP Channels: What Do They Look Like? In: Neurobiology of TRP Channels. 2nd Edition. CRC Press/Taylor & Francis; 2017. doi:10.4324/9781315152837-1

10. Bertl A, Slayman CL. Cation-selective channels in the vacuolar membrane of Saccharomyces: Dependence on calcium, redox state, and voltage. Proc Natl Acad Sci U S A. 1990;87(20):7824–7828. doi:10.1073/pnas.87.20.7824

11. Bertl A, Gradmann D, Slayman CL. Calcium-and voltage-dependent ion channels in Saccharomyces cerevisiae. Philos Trans R Soc Lond B Biol Sci. 1992;338(1283):63–72. doi:10.1098/rstb.1992.0129

12. Zhou XL, Batiza AF, Loukin SH, Palmer CP, Kung C, Saimi Y. The transient receptor potential channel on the yeast vacuole is mechanosensitive. Proc Natl Acad Sci U S A. 2003; 100(12): 7105–7110. doi:10.1073/pnas.1230540100

13. Su Z, Zhou X, Loukin SH, Saimi Y, Kung C. Mechanical force and cytoplasmic Ca2+ activate yeast TRPY1 in parallel. J Membr Biol. 2009;227(3):141–150. doi:10.1007/s00232-009-9153-9

14. Hamamoto S, Mori Y, Yabe I, Uozumi N. In vitro and in vivo characterization of modulation of the vacuolar cation channel TRPY1 from Saccharomyces cerevisiae. FEBS J. 2018;285(6):1146–1161. doi:10.1111/febs.14399

15. Amini M, Wang H, Belkacemi A, et al. Identification of Inhibitory Ca2+ Binding Sites in the Upper Vestibule of the Yeast Vacuolar TRP Channel. iScience. 2019;11:1–12. doi:10.1016/j.isci.2018.11.037

16. Nikolaev YA, Cox CD, Ridone P, et al. Mammalian TRP ion channels are insensitive to membrane stretch. J Cell Sci. 2019. doi:10.1242/jcs.238360

17. Zhang W, Cheng LE, Kittelmann M, et al. Ankyrin Repeats Convey Force to Gate the NOMPC Mechanotransduction Channel. Cell. 2015. doi:10.1016/j.cell.2015.08.024

18. Yan Z, Zhang W, He Y, et al. Drosophila NOMPC is a mechanotransduction channel subunit for gentle-touch sensation. Nature. 2013;493(7431):221–225. doi:10.1038/nature11685

19. Jin P, Bulkley D, Guo Y, et al. Electron cryo-microscopy structure of the mechanotransduction channel NOMPC. Nature. 2017;547(7661):118–122. doi:10.1038/nature22981

20. Su Z, Anishkin A, Kung C, Saimi Y. The core domain as the force sensor of the yeast mechanosensitive TRP channel. J Gen Physiol. 2011;138(6):627–640. doi:10.1085/jgp.201110693

21. Paknejad N, Hite RK. Structural basis for the regulation of inositol trisphosphate receptors by Ca2+ and IP3. Nat Struct Mol Biol. 2018;25(8):660–668. doi:10.1038/s41594-018-0089-6

22. Cao E, Liao M, Cheng Y, Julius D. TRPV1 structures in distinct conformations reveal activation mechanisms. Nature. 2013. doi:10.1038/nature12823

23. Gao Y, Cao E, Julius D, Cheng Y. TRPV1 structures in nanodiscs reveal mechanisms of ligand and lipid action. Nature. 2016;534(7607):347–351. doi:10.1038/nature17964

24. Pumroy RA, Fluck EC, Ahmed T, Moiseenkova-Bell VY. Structural insights into the gating mechanisms of TRPV channels. Cell Calcium. 2020;87. doi:10.1016/j.ceca.2020.102168

25. Hughes TET, Lodowski DT, Huynh KW, et al. Structural basis of TRPV5 channel inhibition by econazole revealed by cryo-EM. Nat Struct Mol Biol. 2018. doi:10.1038/s41594-017-0009-1

26. Winkler PA, Huang Y, Sun W, Du J, Lü W. Electron cryo-microscopy structure of a human TRPM4 channel. Nature. 2017. doi:10.1038/nature24674

27. Tang Q, Guo W, Zheng L, et al. Structure of the receptor-activated human TRPC6 and TRPC3 ion channels. Cell Res. 2018. doi:10.1038/s41422-018-0038-2

28. Nascimbeni AC, Codogno P, Morel E. Phosphatidylinositol-3-phosphate in the regulation of autophagy membrane dynamics. FEBS J. 2017. doi:10.1111/febs.13987

29. Zhou X, Su Z, Anishkin A, et al. Yeast screens show aromatic residues at the end of the sixth helix anchor transient receptor potential channel gate. Proc Natl Acad Sci U S A. 2007. doi:10.1073/pnas.0704039104

30. Zhao J, Lin King J V., Paulsen CE, Cheng Y, Julius D. Irritant-evoked activation and calcium modulation of the TRPA1 receptor. Nature. 2020. doi:10.1038/s41586-020-2480-9

31. Su Z, Zhou X, Haynes WJ, et al. Yeast gain-of-function mutations reveal structure-function relationships conserved among different subfamilies of transient receptor potential channels. Proc Natl Acad Sci U S A. 2007. doi:10.1073/pnas.0708584104

32. Zhao J, Blunck R. The isolated voltage sensing domain of the Shaker potassium channel forms a voltage-gated cation channel. Elife. 2016. doi:10.7554/eLife.18130

33. Geragotelis AD, Wood ML, Göddeke H, et al. Voltage-dependent structural models of the human Hv1 proton channel from long-timescale molecular dynamics simulations. Proc Natl Acad Sci U S A. 2020. doi:10.1073/pnas.1920943117

34. Ramsey IS, Moran MM, Chong JA, Clapham DE. A voltage-gated proton-selective channel lacking the pore domain. Nature. 2006. doi:10.1038/nature04700

35. Sethi A, Eargle J, Black AA, Luthey-Schulten Z. Dynamical networks in tRNA: Protein complexes. Proc Natl Acad Sci U S A. 2009. doi:10.1073/pnas.0810961106

36. Kung C et al. Sensing with Ion Channels (ed Boris Martinac). Springer Berlin Heidelb. 2008:1–23.

37. Hilton JK, Kim M, Van Horn WD. Structural and Evolutionary Insights Point to Allosteric Regulation of TRP Ion Channels. Acc Chem Res. 2019;52(6):1643–1652. doi:10.1021/acs.accounts.9b00075

38. Palmer CP, Zhou XL, Lin J, Loukin SH, Kung C, Saimi Y. A TRP homolog in Saccharomyces cerevisiae forms an intracellular Ca2+-permeable channel in the yeast vacuolar membrane. Proc Natl Acad Sci U S A. 2001. doi:10.1073/pnas.141036198

39. Huffer KE, Aleksandrova AA, Jara-Oseguera A, Forrest LR, Swartz KJ. Global alignment and assessment of TRP channel transmembrane domain structures to explore functional mechanisms. bioRxiv. January 2020:2020.05.14.096792. doi:10.1101/2020.05.14.096792

40. Illergård K, Ardell DH, Elofsson A. Structure is three to ten times more conserved than sequence—A study of structural response in protein cores. Proteins Struct Funct Bioinforma. 2009;77(3):499–508. doi:10.1002/prot.22458

41. McGoldrick LL, Singh AK, Demirkhanyan L, et al. Structure of the thermo-sensitive TRP channel TRP1 from the alga Chlamydomonas reinhardtii. Nat Commun. 2019. doi:10.1038/s41467-019-12121-9

42. Teng J, Loukin S, Anishkin A, Kung C. The force-from-lipid (FFL) principle of mechanosensitivity, at large and in elements. Pflugers Arch. 2015;467(1):27–37. doi:10.1007/s00424-014-1530-2

43. Autzen HE, Myasnikov AG, Campbell MG, Asarnow D, Julius D, Cheng Y. Structure of the human TRPM4 ion channel in a lipid nanodisc. Science (80-). 2018;359(6372):228–232. doi:10.1126/science.aar4510

44. Diver MM, Cheng Y, Julius D. Structural insights into TRPM8 inhibition and desensitization. Science (80-). 2019. doi:10.1126/science.aax6672

45. Dunn T, Gable K, Beeler T. Regulation of cellular Ca2+ by yeast vacuoles. J Biol Chem. 1994;269(10):7273–7278. http://www.jbc.org/content/269/10/7273.abstract.

46. Gordienko D V., Bolton TB. Crosstalk between ryanodine receptors and IP3 receptors as a factor shaping spontaneous Ca2+-release events in rabbit portal vein myocytes. J Physiol. 2002. doi:10.1113/jphysiol.2001.015966

47. Cruickshank CC, Minchin RF, Le Dain AC, Martinac B. Estimation of the pore size of the large-conductance mechanosensitive ion channel of Escherichia coli. Biophys J. 1997;73(4):1925–1931. doi:10.1016/S0006-3495(97)78223-7

48. Hille B. Ion Channels of Excitable Membranes. 3rd ed. Sinauer Associates is an imprint of Oxford University Press; 2001.

49. Moiseenkova-Bell VY, Stanciu LA, Serysheva II, Tobe BJ, Wensel TG. Structure of TRPV1 channel revealed by electron cryomicroscopy. Proc Natl Acad Sci U S A. 2008. doi:10.1073/pnas.0711835105

50. Zivanov J, Nakane T, Forsberg BO, et al. New tools for automated high-resolution cryo-EM structure determination in RELION-3. Egelman EH, Kuriyan J, eds. Elife. 2018;7:e42166. doi:10.7554/eLife.42166

51. Scheres SHW. Processing of Structurally Heterogeneous Cryo-EM Data in RELION. In: Crowther RABT-M in E, ed. Methods in Enzymology. Vol 579. Academic Press; 2016:125–157. doi:10.1016/bs.mie.2016.04.012

52. Scheres SHW. RELION: Implementation of a Bayesian approach to cryo-EM structure determination. J Struct Biol. 2012;180(3):519–530. doi:10.1016/j.jsb.2012.09.006

53. Zheng SQ, Palovcak E, Armache JP, Verba KA, Cheng Y, Agard DA. MotionCor2: Anisotropic correction of beam-induced motion for improved cryo-electron microscopy. Nat Methods. 2017;14(4):331–332. doi:10.1038/nmeth.4193

54. Rohou A, Grigorieff N. CTFFIND4: Fast and accurate defocus estimation from electron micrographs. J Struct Biol. 2015;192(2):216–221. doi:10.1016/j.jsb.2015.08.008

55. Yang J, Zhang Y. I-TASSER server: New development for protein structure and function predictions. Nucleic Acids Res. 2015;43(W1):W174–W181. doi:10.1093/nar/gkv342

56. Roy A, Kucukural A, Zhang Y. I-TASSER: A unified platform for automated protein structure and function prediction. Nat Protoc. 2010;5(4):725–738. doi:10.1038/nprot.2010.5

57. Emsley P, Cowtan K. Coot: Model-building tools for molecular graphics. Acta Crystallogr Sect D Biol Crystallogr. 2004;60(12 I):2126–2132. doi:10.1107/S0907444904019158

58. Moriarty NW, Grosse-Kunstleve RW, Adams PD. Electronic ligand builder and optimization workbench (eLBOW): A tool for ligand coordinate and restraint generation. Acta Crystallogr Sect D Biol Crystallogr. 2009;65(10):1074–1080. doi:10.1107/S0907444909029436

59. Adams PD, Ralf W, Read RJ, et al. PHENIX: Building new software for automated crystallographic structure determination. Acta Crystallogr Sect D Biol Crystallogr. 2002;58(11):1948–1954. doi:10.1107/S0907444902016657

60. Chen VB, Arendall WB, Headd JJ, et al. MolProbity: All-atom structure validation for macromolecular crystallography. Acta Crystallogr Sect D Biol Crystallogr. 2010;66(1):12–21. doi:10.1107/S0907444909042073

61. Barad BA, Echols N, Wang RYR, et al. EMRinger: Side chain-directed model and map validation for 3D cryo-electron microscopy. Nat Methods. 2015;12(10):943–946. doi:10.1038/nmeth.3541

62. Pettersen EF et al. UCSF Chimera--a visualization system for exploratory research and analysis. J Comput Chem. 2004;25:1605–1612. doi:doi:10.1002/jcc.20084

63. Bell JM, Chen M, Baldwin PR, Ludtke SJ. High resolution single particle refinement in EMAN2.1. Methods. 2016;100:25–34. doi:10.1016/j.ymeth.2016.02.018

64. Smart OS, Neduvelil JG, Wang X, Wallace BA, Sansom MSP. HOLE: A program for the analysis of the pore dimensions of ion channel structural models. J Mol Graph. 1996;14(6):354–360. doi:10.1016/S0263-7855(97)00009-X

65. Dolinsky TJ, Nielsen JE, McCammon JA, Baker NA. PDB2PQR: An automated pipeline for the setup of Poisson-Boltzmann electrostatics calculations. Nucleic Acids Res. 2004;32(WEB SERVER ISS.):W665–W667. doi:10.1093/nar/gkh381

66. Schrodinger L. The PyMOL Molecular Graphics System, Version 1.8. 2015.

67. Phillips JC, Braun R, Wang W, et al. Scalable molecular dynamics with NAMD. J Comput Chem. 2005. doi:10.1002/jcc.20289

68. Phillips JC, Hardy DJ, Maia JDC, et al. Scalable molecular dynamics on CPU and GPU architectures with NAMD. J Chem Phys. 2020. doi:10.1063/5.0014475

69. Kelley LA, Mezulis S, Yates CM, Wass MN, Sternberg MJE. The Phyre2 web portal for protein modeling, prediction and analysis. Nat Protoc. 2015. doi:10.1038/nprot.2015.053

70. Wang Q, Corey RA, Hedger G, et al. Lipid Interactions of a Ciliary Membrane TRP Channel: Simulation and Structural Studies of Polycystin-2. Structure. 2020;28(2):169–184.e5. doi:https://doi.org/10.1016/j.str.2019.11.005

71. Sakipov S, Sobolevsky AI, Kurnikova MG. Ion Permeation Mechanism in Epithelial Calcium Channel TRVP6. Sci Rep. 2018. doi:10.1038/s41598-018-23972-5

72. Humphrey W, Dalke A, Schulten K. VMD: Visual molecular dynamics. J Mol Graph. 1996. doi:10.1016/0263-7855(96)00018-5

73. Vanommeslaeghe K, Mackerell AD. CHARMM additive and polarizable force fields for biophysics and computer-aided drug design. Biochim Biophys Acta-Gen Subj. 2015. doi:10.1016/j.bbagen.2014.08.004

74. Jorgensen WL, Chandrasekhar J, Madura JD, Impey RW, Klein ML. Comparison of simple potential functions for simulating liquid water. J Chem Phys. 1983. doi:10.1063/1.445869

75. Glykos NM. Carma: A Molecular Dynamics Analysis Program. J Comput Chem. 2006;27(14): 1765–1768. doi:10.1002/jcc

